# Pattern Recognition Receptor for Bacterial Lipopolysaccharide in the Cytosol of Human Macrophages

**DOI:** 10.1101/2021.10.22.465470

**Authors:** Maricarmen Rojas-Lopez, Amanda S. Zajac, Thomas E. Wood, Kelly A. Miller, María Luisa Gil-Marqués, Austin C. Hachey, Vritti Kharbanda, Keith T. Egger, Marcia B. Goldberg

**Affiliations:** Center for Bacterial Pathogenesis, Division of Infectious Diseases, Massachusetts General Hospital, Boston, MA, USA; Department of Microbiology, Blavatnik Institute, Harvard Medical School, Boston, MA, USA; Broad Institute of MIT and Harvard, Cambridge, MA, USA; Department of Immunology and Infectious Diseases, Harvard T.H. Chan School of Public Health, Boston, MA, USA

## Abstract

Endotoxin - bacterial lipopolysaccharide (LPS) - is a driver of the lethal infection sepsis through activation of innate immune responses. When delivered to the cytosol of macrophages, LPS (cLPS) induces the assembly of an inflammasome that contains caspases-4/5 in humans or caspase-11 in mice. Whereas activation of all other inflammasomes is triggered by sensing of pathogen products by a specific host cytosolic pattern recognition receptor protein, whether pattern recognition receptors for cLPS exist has been doubted by many investigators, as caspases-4, -5, and -11 bind and activate LPS directly *in vitro*. Here we show that the primate-specific protein NLRP11 is a pattern recognition receptor for cLPS required for efficient activation of the caspase-4 inflammasome in human macrophages. *NLRP11* is present in humans and other primates, but absent in mice, likely explaining why it has been missed in screens looking for innate immune signaling molecules, most of which have been carried out in mice. NLRP11 is a previously missing link and a component of the human caspase-4 inflammasome activation pathway.

**One Sentence Summary:** Discovery that human macrophages contain a cytosolic receptor for bacterial lipopolysaccharide.

## INTRODUCTION

Inflammasomes assemble in response to the presence in the cytosol of pathogen-associated molecules, including dsDNA, flagellin, type 3 secretion system structural proteins, pore-forming toxins, ion flux, ATP, and LPS (*1-5*). Canonical inflammasomes consist of a pattern recognition receptor, which senses a pathogen-associate molecule protein, and an inflammatory caspase, frequently linked by an adaptor protein. Inflammasome assembly activates the caspase, which proteolytically activates the pore-forming protein gasdermin D to form plasma membrane pores, leading to pyroptotic cell death (*6-10*). The presence of bacterial LPS in the cytosol of human macrophages activates a non-canonical inflammasome, which in humans, contains caspases-4 and -5, and in mice, caspase-11 (*11-13*), and which also leads to caspase-mediated processing of gasdermin D, pore formation, and pyroptotic cell death. To date, no pattern recognition receptor protein of cytosolic LPS (cLPS) has been identified, and because caspase-4, -5, and -11 bind LPS *in vitro* (*14*), the current paradigm is that the non-canonical inflammasome lacks a pattern recognition receptor.

Here we identify primate specific nucleotide-binding oligomerization domain, leucine rich repeat and pyrin domain containing protein-11 (NLRP11) as a cytosolic pattern recognition receptor of bacterial LPS in human macrophages. We show that efficient activation of the non-canonical pathway in human-derived macrophages depends on NLRP11, and that NLRP11 interacts with core components of the caspase-4 inflammasome. Our results demonstrate that NLRP11 senses cLPS and promotes LPS-dependent activation of caspase-4.

## RESULTS

### NLRP11 required for cell death due to intracellular gram-negative pathogens

In a forward genetic screen that enriched for genes required for death during infection with the gram-negative intracellular bacterial pathogen *Shigella flexneri*, we identified NLRP11 (*15*) (Fig. 1A, and fig. S1); the screen was conducted using a human-derived myeloid lineage cell line. The NLRP family includes NLRP1 and NLRP3, which sense a range of pathogen-associated molecular patterns in the cell cytosol. In human-derived monocyte THP-1 cells containing either of two targeted deletions of *nlrp11*, each homozygous for the indicated deletion (fig. S2), cell death (release of lactate dehydrogenase, LDH) during *S. flexneri* infection was decreased by >50% (Fig. 1B). The efficiency of cell invasion by *S. flexneri* was unaffected by the absence of NLRP11 (fig. S3A), and *S. flexneri* mutants unable to access the cytosol by virtue of inactivation of the type 3 secretion system induced minimal cell death, indicating that cell death depended on bacterial cytosolic localization and/or type 3 secretion activity (fig. S3B).

**Fig. 1.**
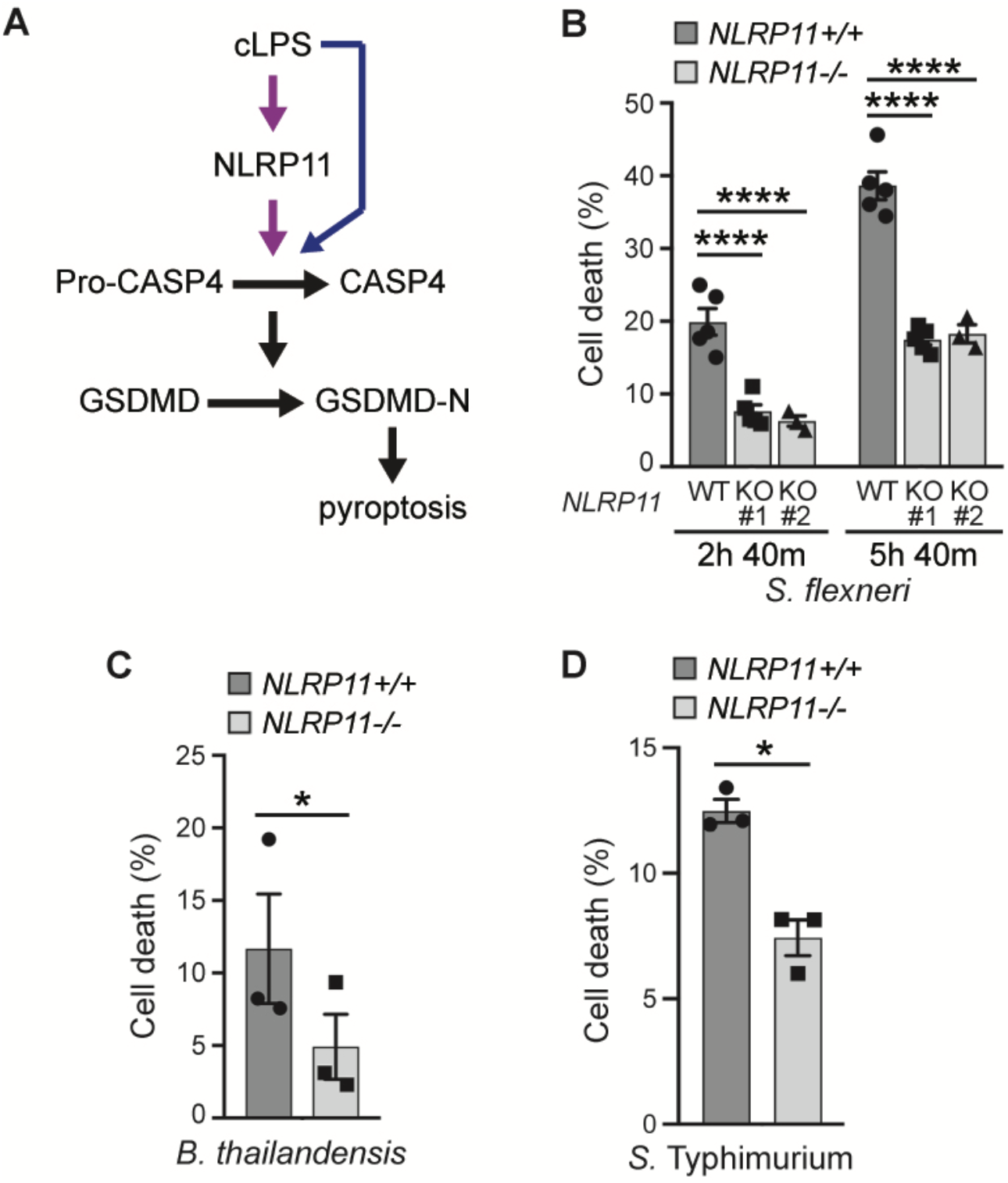
*NLRP11* is required for efficient cell death induced by infection of human-derived macrophage-like cells with intracellular gram-negative bacteria. (**A**) Prevailing (blue/black arrows) and proposed (pink/black arrows) paradigms of non-canonical recognition of cLPS, high cLPS concentration activating both pathways. CASP4, caspase-4; GSDMD, gasdermin D; GSDMD-N, GSDMD N-terminal domain. (**B**-**C**) *NLRP11*-dependent death of THP-1 macrophages upon infection with cytosolic *S. flexneri* (**B**) or *B. thailandensis* (**C**), or predominantly vacuolar *S*. Typhimurium (**D**), in two independent (**B**) or one (**C-D**) *NLRP11*^−/-^ THP-1 cell lines. *n* = 5 (**B**: WT, KO #1) or 3 experiments (**B**: KO #2, **C-D**). Cell death, by LDH release, as percent of Triton X-100 cell lysis. Mean ± S.E.M. Two-way ANOVA with Dunnett *post hoc* test (**B, D**), or student T-test (**C**). *, p<0.05; ***; p<0.001; ****, p<0.0001.

To test whether the dependence on NLRP11 for efficient cell death was generalizable to other intracellular gram-negative bacterial pathogens, we infected *NLRP11*^−/-^ cells with the cytosolic pathogen *Burkholderia thailandensis* and the largely vacuolar pathogen *Salmonella enterica* serovar Typhimurium. As with *S. flexneri*, in *NLRP11*^−/-^ macrophages, cell death during infection with *B. thailandensis* or *S*. Typhimurium was significantly reduced (Fig. 1, C and D). Upon viral infection and certain Toll-like receptor interactions, the NF-κB pathway is repressed by NLRP11-dependent targeted degradation of TRAF6 (*16-18*); however, *S. flexneri*-induced NLRP11-dependent cell death was independent of TRAF6, whether or not *Shigella* OspI, which suppresses TRAF6 signaling (*19*), was present (fig. S3, C, D, and E), indicating that the role of NLRP11 in bacterial infection-induced cell death is distinct from its role in NF-κB signaling. The levels of most other innate immune proteins examined were unchanged by disruption of *NLRP11* (fig. S4), with the exception that levels of NLRP3 were increased in *NLRP11*^−/-^ cells, likely a result of TRAF6-dependent derepression of NF-κB activation of *NLRP3* expression. Moreover, tyrosine phosphorylation of STAT1 in response to IFNβ was only slightly diminished by the absence of NLRP11 (fig. S5). Thus, NLRP11 is required for efficient cell death induced by multiple intracellular gram-negative pathogens.

### Bacterial LPS recognized by NLRP11 in cytosol

We tested whether LPS, which is specific to gram-negative bacteria, might be required for NLRP11-dependent cell death. Infection of THP-1 macrophages with the cytosolic gram-positive pathogen *Listeria monocytogenes* induced minimal levels of cell death during the experimental timeframe, but when coated with *S. enterica* LPS, induced cell death that was largely dependent on NLRP11 (Fig. 2A). The observed cell death was dependent on cytosolic localization, as *L. monocytogenes* lacking listeriolysin O (LLO), which is required to lyse the vacuole, induced minimal cell death. Introduction of LPS alone directly into the cytosol by electroporation similarly induced cell death that was significantly dependent on NLRP11, both in *NLRP11*^−/-^ and NLRP11 siRNA-treated THP-1 macrophages (Fig. 2, B and C), whereas cell death induced by extracellular LPS was independent of NLRP11 (fig. S6). Therefore, NLRP11 functions in a cell death pathway that is activated by LPS localized to the cytosol (cLPS).

**Fig. 2.**
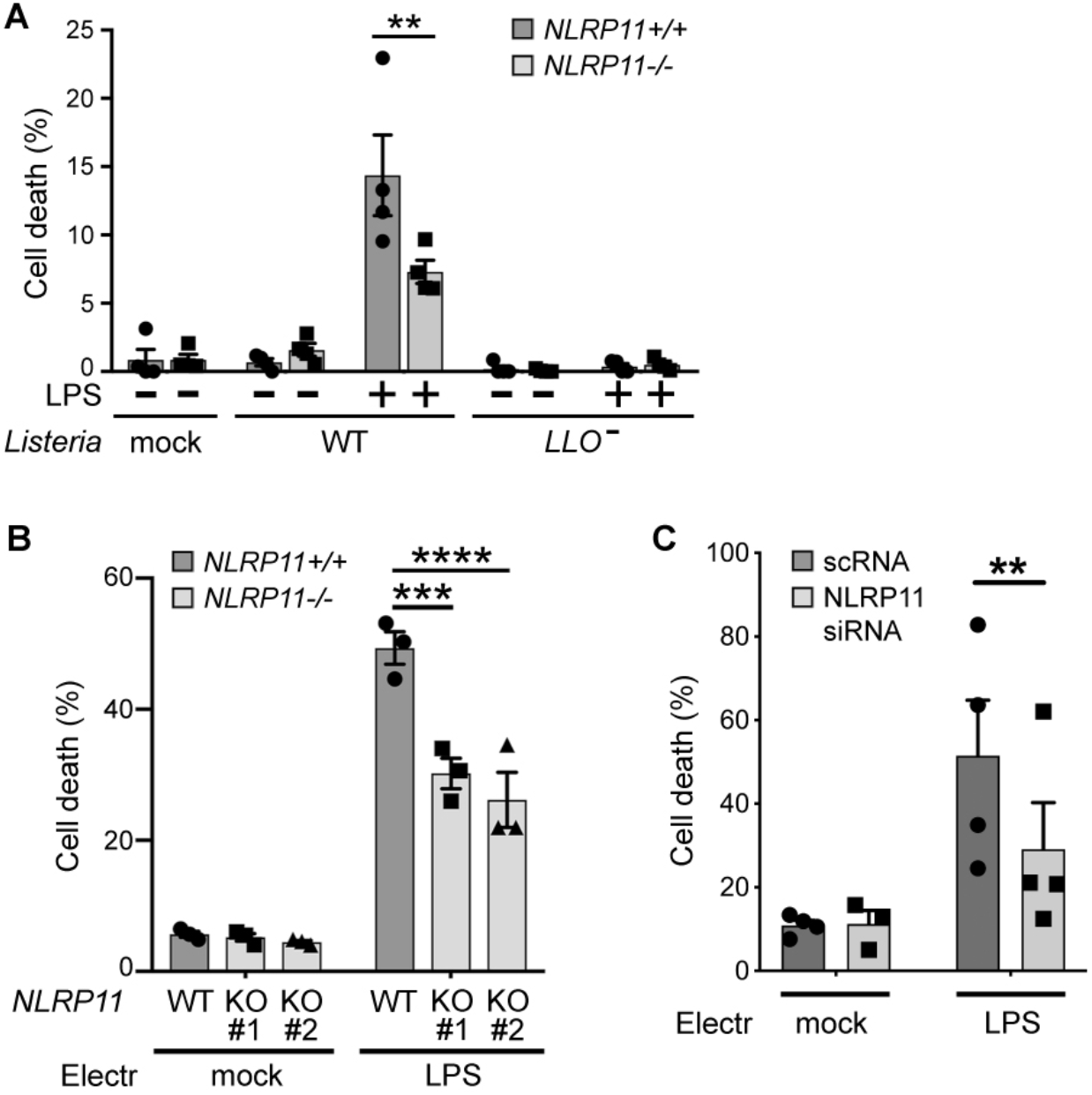
*NLRP11* potentiates macrophage cell death in response to cytosolic LPS. **A**) *NLRP11*-dependent death of THP-1 macrophage-like cells upon infection with cytosolic gram-positive pathogen *L. monocytogenes* coated with *S. enterica* serovar Minnesota LPS. Cell death depends on LPS and vacuolar escape, as *L. monocytogenes LLO*^−^ is restricted to the phagosome. *n* = 4 experiments. (**B-C**) In *NLRP11*^−/-^ THP-1 macrophages (**B**) or following NLRP11 siRNA (**C**), cell death in response to electroporated LPS is defective. *n* = 3 (**B**) or 4 (**C**) experiments. Cell death measured as in Fig 1. Electr, electroporation; mock, carrier alone; scRNA, scrambled siRNA. Two-way ANOVA with Sidak *post hoc* test (**A**) or Tukey *post hoc* test (**B-C**). Mean ± S.E.M. **, p<0.01; ***; p<0.001; ****, p<0.0001.

### NLRP11 is required for efficient activation of non-canonical inflammasome

Activation of caspase-4 was dependent on NLRP11; during *S. flexneri, S*. Typhimurium, or *B. thailandensis* infection, processing of caspase-4, which releases its active domains, was significantly reduced in culture supernatants of *NLRP11*^−/-^ macrophages, yet the amount of pro-caspase-4 in cell lysates was unaltered by disruption of *NLRP11* (Fig. 3, A and B, and fig. S7, A and B). Caspase-1 activation followed a pattern parallel to that of caspase-4, with diminished processing in *NLRP11*^−/-^ macrophages (fig S7C); the pattern of activation of caspase-1 was expected, as K+ efflux that results from activation of the non-canonical inflammasome activates NLRP3 and caspase-1 (*6, 20-22*).

**Fig. 3.**
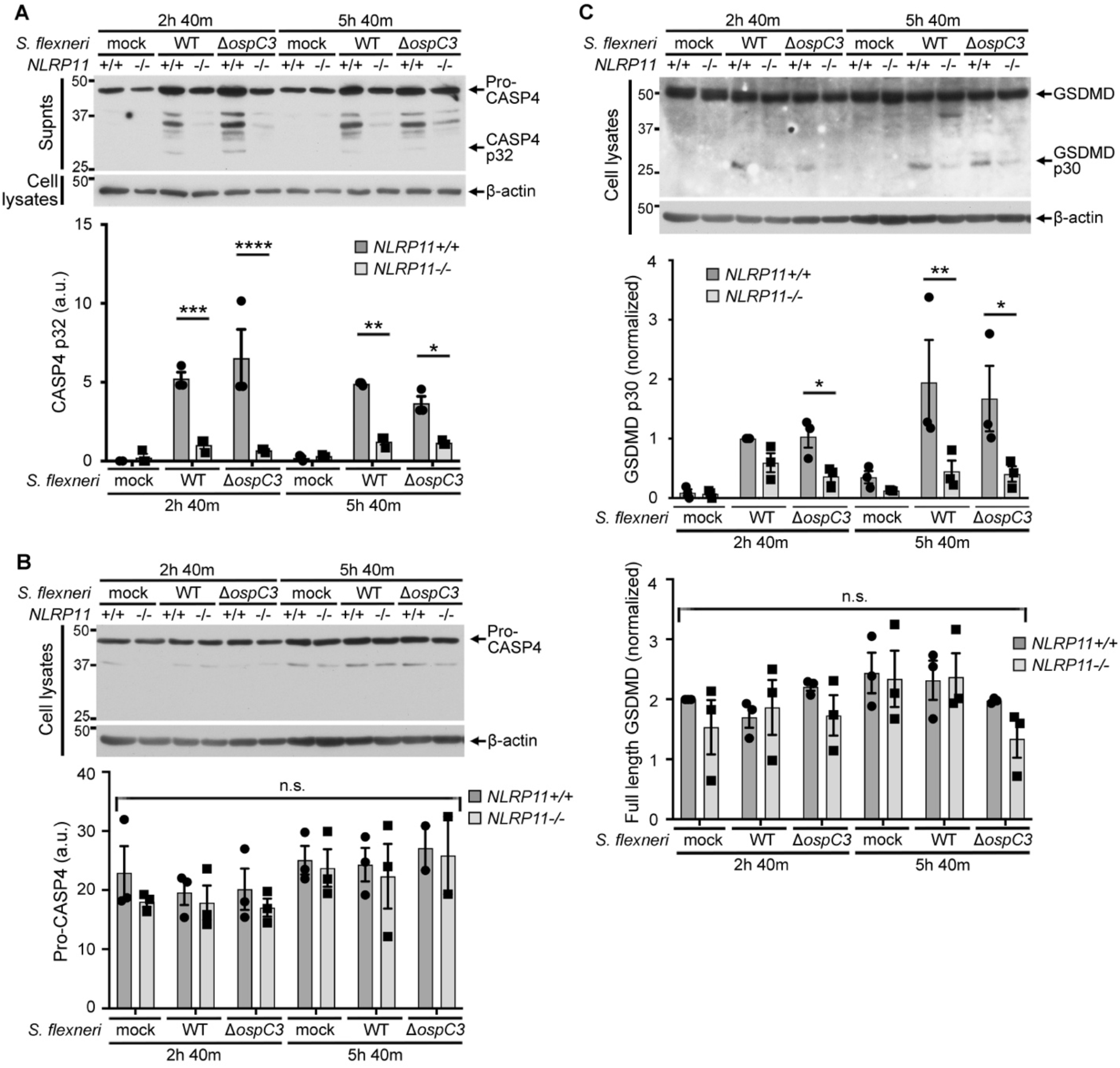
*NLRP11* is required for processing of caspase-4 and gasdermin D during *S. flexneri* infection. *S. flexneri* infection of THP-1 macrophages. (**A**) Processing of caspase-4 (CASP4), with release of p32 polypeptide. Top, representative western blots of culture supernatants (CASP4) and associated cell lysates (β-actin); bottom, corresponding band densitometry of CASP4 p32. (**B**) Levels of pro-CASP4 in cell lysates from experiments described in panel **A**. Top, representative western blots of cell lysates; bottom, corresponding band densitometry of pro-CASP4. (**C**) Processing of gasdermin D (GSDMD), with release of p30 polypeptide. Top, representative western blots of cell lysates; bottom, corresponding band densitometry of GSDMD p30 (top) and full length GSDMD (bottom); in (**C**), densitometry is normalized to band for wildtype (WT) *S. flexneri* infection of NLRP11^+/+^ THP-1 macrophages. In the absence of OspC3, processing was similar to induced by WT *S. flexneri*; in mouse bone marrow-derived macrophages and human epithelial cell lines, OspC3 inhibits caspase-4 processing of gasdermin D (*56, 57*). *n* = 3 experiments. MW markers in kD. Mean ± S.E.M. Two-way ANOVA with Sidak *post hoc* test. *, p<0.05; **, p<0.01; ***; p<0.001; ****, p<0.0001; n.s., not significant for any comparison.

A role of the non-canonical inflammasome in *S. flexneri* induced cell death is consistent with previously published data showing that in mouse macrophages, in the absence of caspase-11, cell death induced by *S. flexneri* infection is reduced by 40-50% (*6, 23*); of note, earlier studies of *S. flexneri*-induced death of murine macrophages must be interpreted with caution, as it was learned in 2011 that the macrophages used in these early studies lacked both caspase-1 and caspase-11 (*11*). Activation of gasdermin D was also significantly reduced in *NLRP11*^−/-^ macrophages, yet levels of full-length gasdermin D in cell lysates were independent of NLRP11 (Fig. 3C and fig. S4, A and C). Moreover, release of the IL-1 family cytokines IL-1β and IL-18 from *S. flexneri*-infected *NLRP11*^−/-^ macrophages was defective (fig. S8).

Upon *CASP4* depletion, the absence of NLRP11 caused no significant additional decrease in cell death induced by LPS electroporation (Fig. 4, A and B). Depletion of *CASP1* induced no significant decrease in cell death in response to cLPS as compared to control depletion for either *NLRP11*^−/-^ or WT THP-1 macrophages (Fig. 4, A and C). These data indicate that NLRP11 is required for efficient activation of the non-canonical inflammasome, induces caspase-4-dependent cell death in the presence of cytosolic LPS, and that NLRP11 function in response to cytosolic LPS is largely independent of caspase-1.

**Fig. 4.**
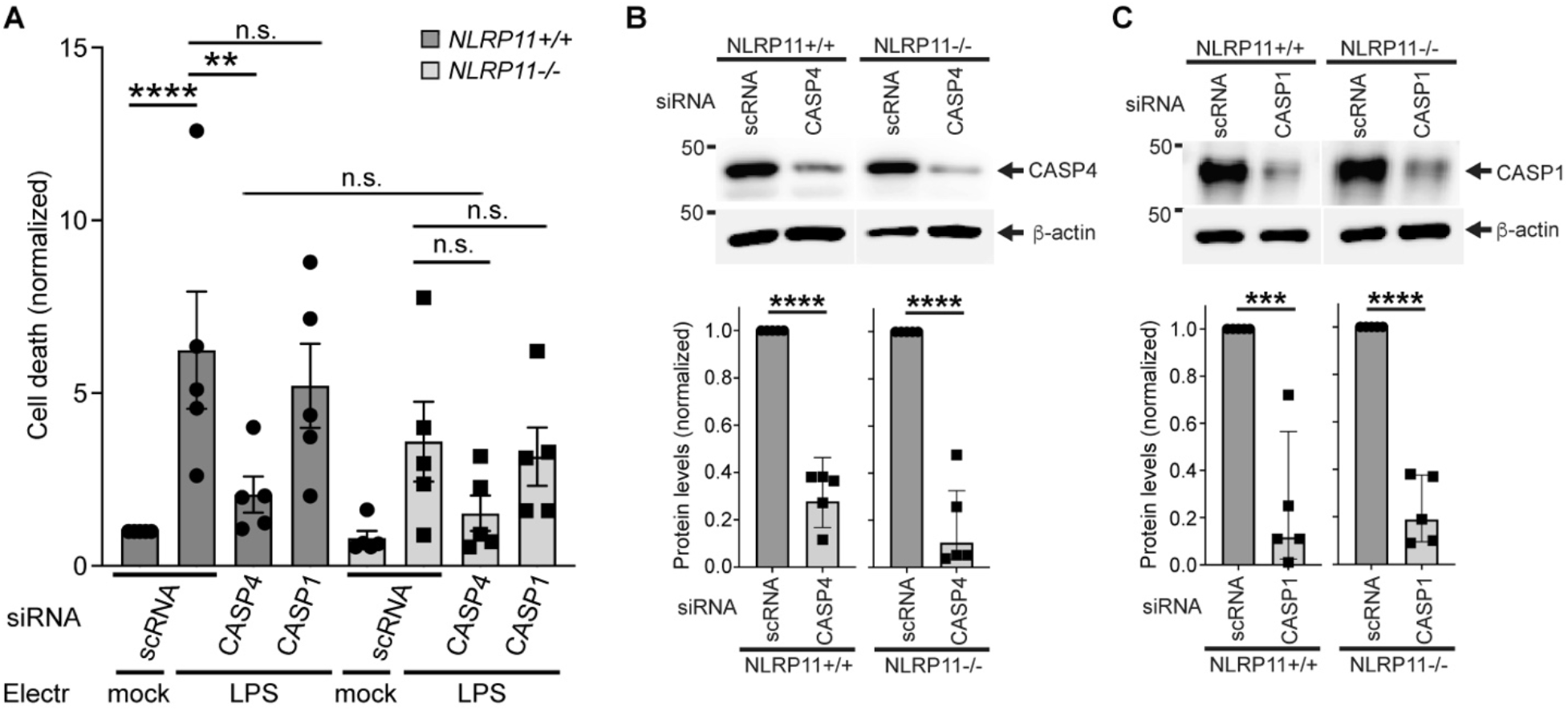
NLRP11-dependent cell death in response to cLPS is dependent on caspase-4 but independent of caspase-1. (**A**) Cell death of THP-1 macrophages in response to electroporated LPS following siRNA depletion of *CASP4* or *CASP1. n* = 5 experiments. Cell death measured as in Fig 1; normalized to mock transfection of *NLRP11*^+/+^ cells following scRNA treatment because data from 5 independent experiments were combined. (**B-C**) Extent of depletion with siRNA of *CASP4* (**B**) or *CASP1* (**C**). *n* = 5 experiments. Representative western blots of cell lysates and corresponding band densitometry for each knocked down protein, normalized to level or protein in scRNA condition. The western blot panels for each protein within each panel are from the same blot. β-actin, loading control. Electr, electroporation; mock, carrier alone; scRNA, scrambled siRNA; CASP4, caspase-4; CASP1, caspase-1. Two-way ANOVA with Tukey *post hoc* test (**A**) or unpaired student T-test (**B-C**). Mean ± S.E.M. **, p<0.01; ***, p<0.001; ****, p<0.0001; n.s., not significant.

### NLRP11 interaction with LPS and caspase-4

In contrast to NLRP3 inflammasomes, activation of the non-canonical pathway is independent of the adaptor protein ASC (*11, 14, 16*). We tested whether NLRP11 interacts with other core components of the non-canonical pathway by ectopic expression of selected components in HEK293T cells, which lack most innate immune proteins, including ASC and caspase-4 (fig. S9). Biotin-conjugated LPS precipitated NLRP11 in the absence of caspase-4 (Fig. 5A), indicating that NLRP11 binds LPS independently of caspase-4. These data do not distinguish whether LPS binding to NLRP11 is direct or indirect. LPS specifically precipitated caspase-4 and its catalytically inactive derivative (CASP4 C258A), as described (*14*), including in the presence of NLRP11 (Fig. 5B), whereas the triacylated lipid Pam3CSK4 did not precipitate either NLRP11 or caspase-4 (Fig. 5A). In addition to binding to LPS independent of caspase-4, NLRP11 specifically and reciprocally precipitated caspase-4 (Fig. 5C and Fig. 6). We developed a plate binding assay using FLAG-NLRP11 in HEK293T lysates and purified caspase-4 (fig. S10). FLAG-NLRP11 binding depended on caspase-4, as in the absence of caspase-4, bound FLAG-NLRP11 was at background levels (Fig. 5D, diamonds). Binding of NLRP11 to caspase-4 increased as a function of increasing concentrations of both NLRP11 lysate and caspase-4, resembling the asymptotic phase of a sigmoidal curve, beginning to reach saturation at 8-10 µg/ml caspase-4 coating the wells (Fig. 5D). Thus, NLRP11 separately binds LPS and caspase-4, and does so independently of ASC.

**Fig. 5.**
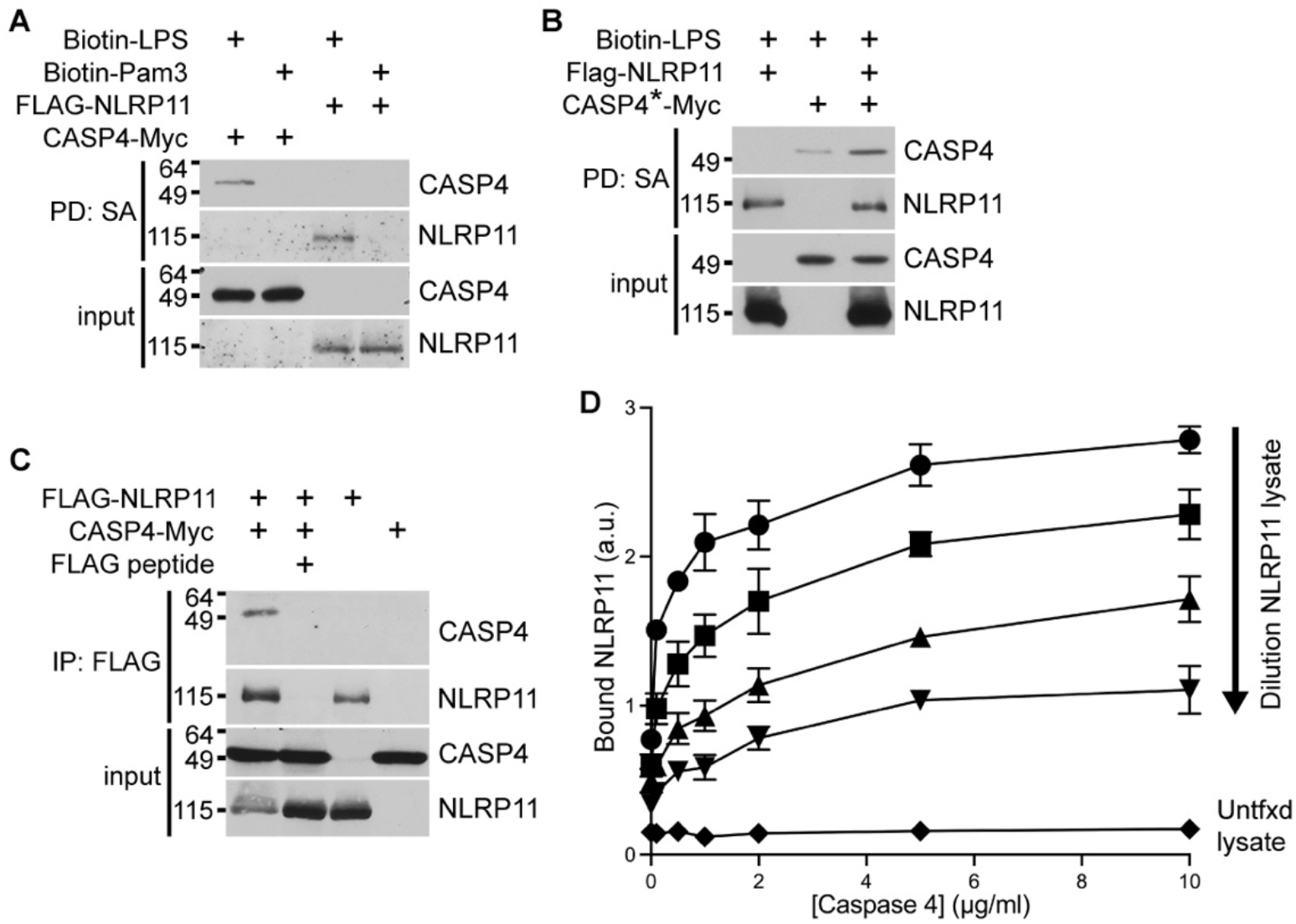
NLRP11 interactions with LPS and caspase-4. Precipitations from reconstituted HEK293T lysates expressing NLRP11 and/or caspase-4. (**A-B**) Precipitation of NLRP11 and wildtype caspase-4 (CASP4) by biotin-LPS, separately (**A**) and together (**B**), and not biotin-Pam3CSK4 (Pam3) (**A**). PD: SA, precipitation with streptavidin beads. *n* = 3 experiments. CASP*, catalytically inactive CASP4 C258A. (**C**) NLRP11 precipitates wildtype CASP4. *n* = 3 experiments. MW markers in kD. (**D**) Increasing interaction between NLRP11 and CASP4 as a function of concentration of each protein. FLAG-NLRP11 in transfected HEK293T cells bound to plates coated with purified wildtype CASP4; detection of bound FLAG. NLRP11 lysate dilutions: circles, undiluted; squares, 1:2 dilution; triangles, 1:4 dilution; inverted triangles, 1:8 dilution. Diamonds, untransfected (untfxd) lysate. *n* = 3 experiments. a.u., arbitrary units.

**Fig. 6.**
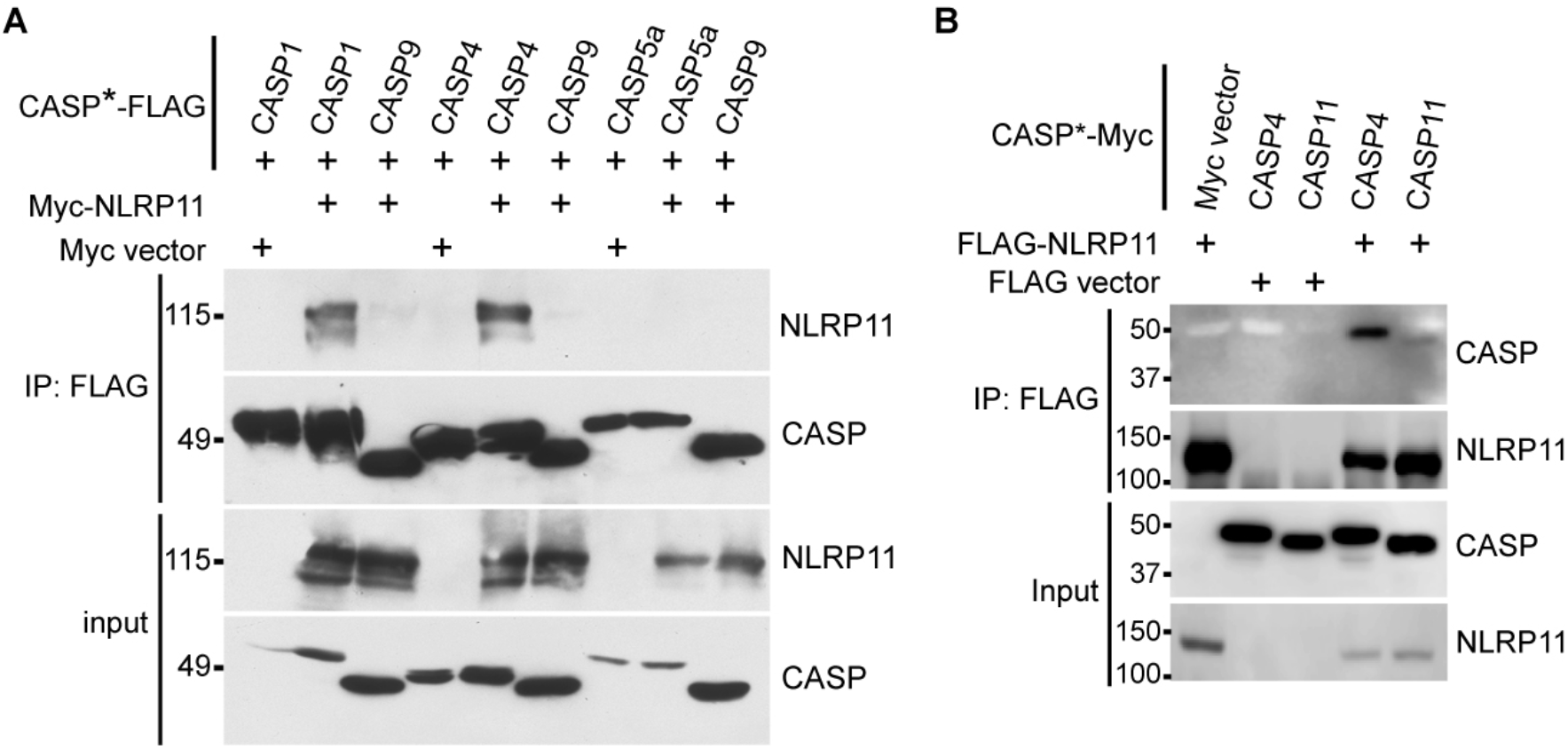
Specificity of NLRP11 for caspases. Precipitations from reconstituted HEK293T lysates expressing NLRP11 and indicated caspases. (**A**) FLAG-tagged inactive caspase (CASP1[C285A]-FLAG, CASP5a[C315A]-FLAG, CASP9[C287S]-FLAG) precipitation of Myc-NLRP11. (**B**) FLAG-tagged NLRP11 precipitation of inactive mouse caspase-11 (CASP11[C254A]-Myc) versus human caspase-4 (CASP4[C258A]-Myc). CASP*, catalytically inactive caspase. IP: FLAG, precipitation with FLAG antibody-coated beads. Representative western blots detection with anti-FLAG or anti-Myc antibody. *n* = 3 experiments (each **A** and **B**). MW markers in kD.

### Caspase specificity of NLRP11

In addition to caspase-4, NLRP11 co-precipitated with caspase-1; it did not co-precipitate with caspase-5, a paralog of caspase-4 that binds LPS and a component of some non-canonical inflammasomes (*14, 24, 25*), with the more divergent human caspase-9, and only minimally with mouse caspase-11, which also binds LPS and is similar but non-identical to human caspase-4 (Fig. 6 and fig. S11). The observed interaction of NLRP11 with caspase-1 may play a role in the recently described contribution of NLRP11 to the NLRP inflammasome (*26*). However, NLRP11 function in cLPS-induced cell death of THP-1 macrophages is independent of caspase-1 (Fig. 4, A and C), indicating that an interaction of caspase-1 with NLRP11 contributes minimally to activation of the non-canonical inflammasome.

### NLRP11 role independent of interferon-gamma and guanylate-binding protein-1

Host GBPs are cytosolic proteins that bind intracellular LPS, promote non-canonical inflammasome activation (*27-31*), and are encoded by genes that are strongly induced by interferon-gamma (IFNγ) (*32*). Human GBP1 binds LPS on the *S. flexneri* surface, whereupon it enhances caspase-4 recruitment (*33*). In the absence of IFNγ pre-treatment, NLRP11-dependent cell death during *S. flexneri* infection appeared to be independent of GBP1, as GBP1 was not induced by *S. flexneri* alone during the experimental timeframe (figs. S4B, and S12, A and B). IFNγ pre-treatment of THP-1 macrophages induced GBP1, as expected, yet even in the presence of GBP1, NLRP11 was required for efficient activation of *S. flexneri*-induced cell death (fig. S12, A and B). These results indicate that in THP-1 macrophages, GBP1 is not required for non-canonical inflammasome activation or pyroptosis or for NLRP11 function in inducing non-canonical pyroptosis. Moreover, IFNγ and GBP1 do not bypass the requirement for NLRP11 in this pathway.

### Requirement for NLRP11 in primary human macrophages

As in THP-1 macrophages, in primary human monocyte-derived macrophages, siRNA depletion of NLRP11 was associated with significantly diminished cLPS-induced cell death (Fig. 7, A and B). Cell death depended on caspase-4 (Fig. 7, C and D), demonstrating that the requirement for NLRP11 in the non-canonical inflammasome pathway extends to primary human cells.

**Fig. 7.**
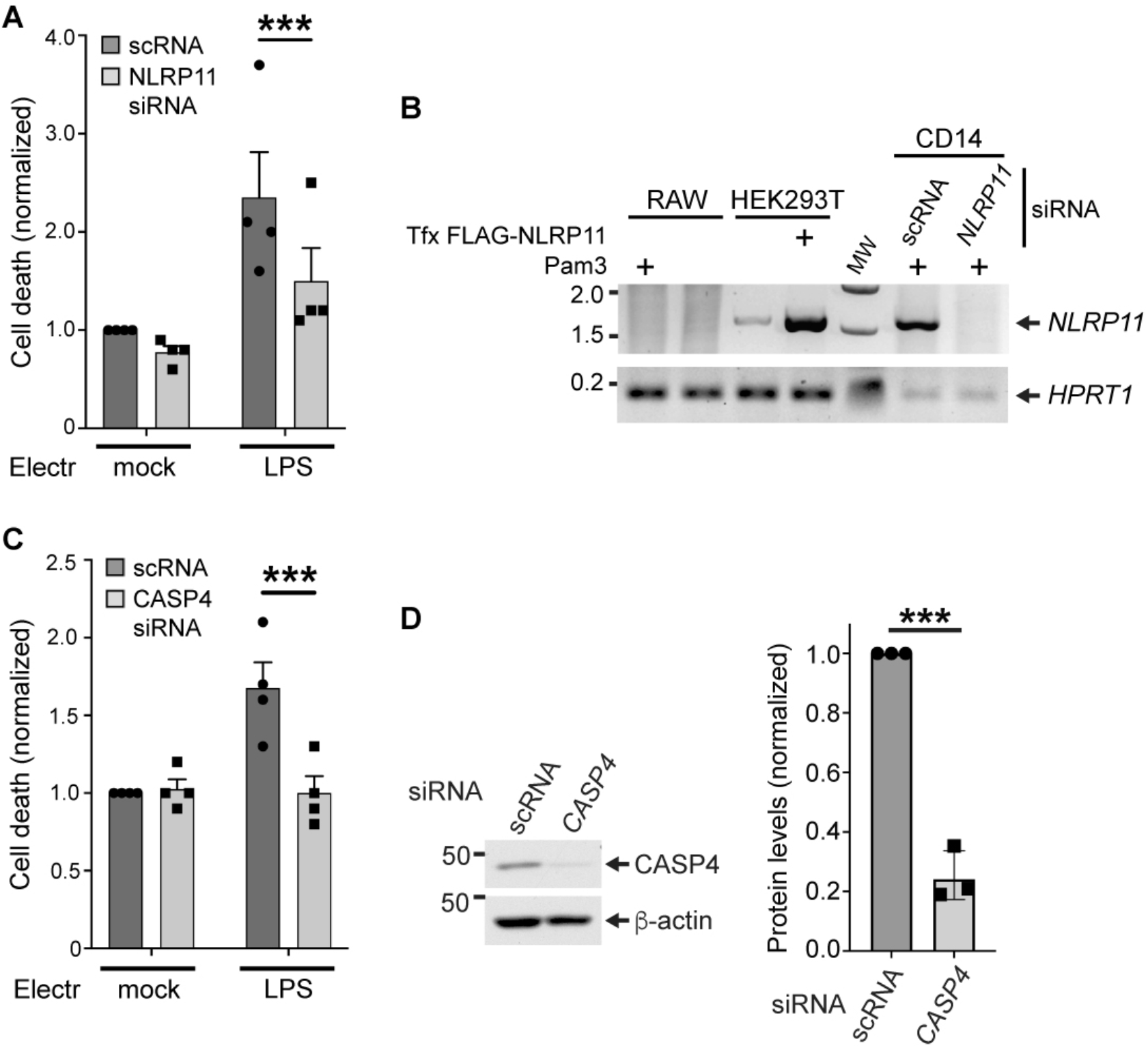
Requirement for NLRP11 for cell death in response to cLPS in primary human macrophages. NLRP11-dependent cell death in response to electroporated LPS in isolated primary human monocyte-derived macrophages (CD14+), treated with siRNA to *NLRP11*. (**A**) Cell death in CD14+ macrophages depleted or not for *NLRP11*. (**B**) RT-PCR analysis of *NLRP11* transcript following siRNA and with or without Pam3 pretreatment. *HRPT1*, housekeeping gene. Representative data from CD14+ cells from a single donor. Negative control, mouse RAW 309 (RAW) macrophages; positive control, HEK293T cells transfected with FLAG-NLRP11 construct. *NLRP11*, PCR amplified fragment of *NLRP11* transcript. (**C**) Cell death in CD14+ macrophages depleted or not for *CASP4*. (**D**) Level of caspase-4 (CASP4) depletion. Representative western blots of cell lysates (left) and corresponding band densitometry, normalized to level or protein in scRNA condition (right). β-actin, loading control. Data are combined from 4 experiments (**A** and **C-D**); each experiment was performed with cells from a different individual donor. Cell death measured as in Fig 1; normalized to mock electroporation following scRNA treatment. *n* = 4 experiments. For cell death measurements, an identical number of cells were used for each condition (see Methods). Electr, electroporation; mock, carrier alone; scRNA, scrambled siRNA. Two-way ANOVA with Šidák’s post-hoc tests (**A, C**); unpaired student T-test (**D**). Mean ± S.E.M. ***; p<0.001.

## DISCUSSION

The current dogma for LPS recognition – that caspase-4 is the sensor for cytosolic LPS – is based on the simplest interpretation of data that show that *in vitro* caspase-4 binds, is induced to oligomerize, and is activated by LPS (*14*). LPS consists of lipid A, an acylated hydrophobic diglucosamine backbone that anchors LPS in the bacterial outer membrane, core oligosaccharide, and serotype-specific O-antigen. The moiety of LPS that binds caspase-4 is lipid A (*14*). We speculate that caspase-4 binding and activation by LPS under the described *in vitro* conditions (*14*) is due, at least in part, to the presence of a multiligand platform of lipid A presented by LPS in micelles or aggregates. Like other caspases, activation of caspase-4 is triggered by proximity-induced dimerization (*14, 34-41*); dimerization is sufficient to activate the caspase to undergo auto-processing, thereby releasing the small and large active subunits, which form an active heterotetrametric complex. As LPS species self-associate in micelles and aggregates at concentrations above critical micelle concentrations, estimated to be 40 nM or lower (*42*), and caspase-4 binds preferentially to LPS in micelles or bacterial outer membrane vesicles (*43*), it seems likely that LPS micelles and aggregates provide a multiligand platform for caspase-4 dimerization. Supporting this possibility is that although at 100-250 nM, purified LPS binds, oligomerizes, and activates caspase-4 *in vitro*, in the presence of >0.25% Tween-20, LPS-mediated caspase-4 oligomerization and activation are disrupted (*14*).

The data presented here identify NLRP11 as a primate-specific pattern recognition receptor of bacterial LPS in the cytosol of human macrophages. As such, NLRP11 is a critical component of activation of the caspase-4 dependent inflammasome and a critical mediator of caspase-4-dependent pyroptosis. Nevertheless, it is likely that delivery of high concentrations of LPS to the macrophage cytosol is sufficient to activate caspase-4 directly and independently of NLRP11, perhaps principally when multiligand LPS platforms are present. Consistent with this, a proportion of caspase-4-dependent cell death appeared to be independent of NLRP11 (Fig. 4, A and B); in these experiments, LPS was electroporated at relatively high concentration, and cells were infected with ten bacteria per cell. We speculate that NLRP11 may be particularly critical for non-canonical inflammasome activation when LPS is delivered in low amounts, such as during human infection with gram-negative pathogens, wherein the intracellular burden of organisms and intracellular LPS is likely very low.

Limitations of the investigations presented here are that, although performed with human cells, they were *ex vivo*, and that due to the restriction of NLRP11 to primates, no mouse model for testing the relevance of NLRP11 in animals currently exists. It remains uncertain whether the unique presence of NLRP11 in primates and the absence of identifiable homologs in mice contributes to the exquisite sensitivity of humans to endotoxic shock as compared with mice; humans display ∼10^4^-fold increased susceptibility to gram-negative bacterial sepsis and bloodstream LPS relative to mice (*44, 45*). Additional insights into these relationships would facilitate much-needed improved mouse models for the study of bacterial sepsis.

## MATERIALS AND METHODS

### Bacterial strains, plasmids, antibodies, and reagents

*Shigella flexneri* serotype 2a wildtype strain 2457T and the *S. flexneri* mutant strains *virB*::Tn5, BS103, and Δ*ospC3*, which are isogenic to 2457T, have been described previously (*46-49*). *Salmonella enterica* serovar Typhimurium and the isogenic Δ*sifA* mutant (*50*) were gifts from L. Knodler. *Burkholderia thailandensis* wild-type strain E264 (ATCC 700388) (*51*) was a gift from P. Cotter. *Listeria monocytogenes* 10403s and 10403 LLO^−^ (*52*) were gifts from M. Kielian and J. Miller. *S. flexneri* was grown overnight in tryptic soy broth at 37°C with aeration. *L. monocytogenes* was grown overnight in brain heart infusion broth at 30°C at a stationary slant. *B. thailandensis* was grown overnight in low salt Luria broth (LB) at 37°C with aeration. *S. enterica* Typhimurium was grown in LB broth at 37°C with aeration.

Caspase-1 (Abcam, ab207802) rabbit monoclonal antibody was used at 0.5 μg/mL (1:1000). Caspase-4 (Santa Cruz, sc-56056), and gasdermin D (Santa Cruz, sc-81868) mouse monoclonal antibodies were used at 0.5 μg/mL (1:200). FLAG M2 mouse monoclonal antibody (Sigma, F3165) was used at 76 ng/mL (1:5000). Myc rabbit monoclonal antibody (Cell Signaling, 2278) was used at 1:1000. GBP1 mouse monoclonal antibody (Abcam, ab119236) was used at 1:1000. Rabbit STAT1 antibody (Cell Signaling Technology, 9172) was used at 32 ng/mL, and rabbit phospho-STAT1 (Tyr-701) antibody (Cell Signaling Technology, 9167) was used at 59 ng/mL.

For western blots of immunoprecipitations, the secondary antibody used was TrueBlot horseradish peroxidase-conjugated anti-mouse Ig (Rockland, 18-8817-31). For detecting β-actin, horseradish peroxidase-conjugated anti-β-actin antibody (Sigma, A3854) was used. For all other western blots, the secondary antibody used was horseradish peroxidase-conjugated goat anti-rabbit or goat anti-mouse IgG antibodies (Jackson ImmunoResearch, 115-035-003 or 111-035-144). All antibodies were diluted in 5% milk in TBST (0.1% Tween).

### Strain and plasmid construction

The *S. flexneri ospI* deletion mutant was generated in the 2457T strain background using the lambda red method by introducing a chloramphenicol acetyltransferase (*cat*) cassette into the *orf169b* gene (which encodes OspI), as described (*53*).

pCMV6-CASP4-MYC and pCMV6-CASP5-MYC were made from pCMV6-CASP4-MYC-FLAG and pCMV6-CASP5-MYC-FLAG, gifts from S. Shin. The isoform of caspase-5 that was used was caspase-5/a, the most abundant isoform; this isoform contains nearly all sequences present in other isoforms. pCMV6-CASP4-MYC C258A and pCMV6-CASP4-MYC K19E C258A were generated by splicing by overlap extension PCR (SOE-PCR). pCMV6-CASP1-MYC C285A and pCMV6-mCASP11-MYC C254A were made from pFastBac-CASP1 C285A (gift from S. Shin) and pMSCV-mCASP11 C254A (gift from J. Kagan), respectively. pCMV6-CASP9-MYC C287S was generated by SOEing PCR from the template pET28a-CASP9 (gift from H. Wu). pcDNA-NLRP11 (gift from H. Wu) was subcloned into pCMV-FLAG with an N-terminal FLAG tag for FLAG-NLRP11 and an N-terminal MYC tag for MYC-NLRP11. All FLAG tags are 3xFLAG. All N-terminal Myc tags are 6xMyc; all C-terminal Myc tags are 1xMyc. All DNA used for transfection was prepared LPS-free (Macherey-Nagel, 740410.50).

### Generation of CRISPR knockouts

THP-1 cells were transduced with lentivirus packaged with pLentiCas9-Blast (Addgene, 52962) generated from HEK293T cells. THP-1 cell suspensions were treated with 8 μg/mL polybrene, combined with lentivirus and centrifuged in 12-well plates for 2 h at 1000 x g at 30°C. After centrifugation, cells were incubated for 2 h and washed once with fresh media. After 48 h, blasticidin (10 μg/mL) was added to the transduced cells. Guide RNA sequence (“sgRNA3”, AGGCATGCAAAGCTGTCATG) was subcloned into pLentiGuide-Puro (Addgene, 59702). Guide RNA expressing plasmids were packaged into lentivirus using HEK293T cells. Polybrene treated THP-1 cells were transduced and selected with 1 μg/mL puromycin. Cells were limiting diluted and single cell clones were screened by SURVEYOR (IDT, 706020) and Sanger sequencing coupled with TIDE analysis (*54*).

### Generation of *TRAF6* depletion cell lines

pLKO.1-shTRAF6 ((*55*), gift of A. Goldfeld), which encodes shRNA to *TRAF6*, was packaged into lentivirus in HEK293T cells. Lentivirus were introduced into NLRP11+/+ and NLRP11-/-THP-1 cell lines and cells containing pLKO.1-shTRAF6 were selected, expanded by maintenance in 0.8 µg/mL puromycin, as described (*55*), and stable clones were isolated by limiting dilution.

### Mammalian cell culture

All cell lines were obtained from ATCC. HEK293T cells were maintained in Dulbecco’s Modified Eagle’s medium (Gibco, 11965118). THP1 cells were maintained in RPMI-1640 (Gibco, 21870092). All media was supplemented with 10% (vol/vol) fetal bovine serum (heat inactivated, sterile filtered, R&D Systems). For THP-1 cells, RPMI-1640 media was also supplemented with 10 mM HEPES (Gibco,15630080) and 2 mM L-glutamine (Gibco, 25030081). All cells were grown at 37°C in 5% CO_2_. To differentiate monocyte cell lines into macrophage-like cells, 50 ng/mL phorbol-12-myristrate-13-acetate (PMA) was added 48 h before experiments. Where indicated, cells were pre-treated with 2.5 or 25 U/mL IFNβ (STEMCELL Technologies, 78113) for 2 h before analysis or 10 ng/mL IFNγ (Millipore, Sigma-Aldrich, I17001) for 16-20 h prior to infection.

### Precipitation experiments

HEK293T cells were transiently transfected using Fugene6 (Promega). A stock solution of 10x Buffer A (100 mM HEPES, 1500 mM NaCl, 10 mM EGTA, 1 mM MgCl_2_, pH 7.4) was prepared. The cell culture medium was removed 24-48 h after transfection. Cells were washed with cold PBS and lysed in 1x Buffer A, 0.5% Triton-X. Cell lysates were incubated on ice for 15 min, vortexed for 15 s and centrifuged at 13,200 rpm for 10 min at 4°C. The pellet was discarded, and Bradford assay was used to determine protein concentration. 30 μL of FLAG M2 Magnetic Beads (Millipore-Sigma, M8823) were washed twice in 1x Buffer A, 0.1% Triton-X and equilibrated in the same buffer at 4°C for 2 h. 500 μg of protein lysate was incubated with the beads for 2 h at 4°C. Beads were washed 3 times with 1x Buffer A, 0.1% Triton-X and eluted with 50 μL of 2x Laemmli buffer.

Streptavidin pulldown was largely as described (*14*), with minor modifications. Cell lysates were diluted to 4 mg/mL before a 2 h incubation with beads. For co-precipitation experiments, separate cells were transfected with each DNA construct, and pre-specified amounts of lysates were combined with each other or with untransfected lysate, such that the total amount of lysate were similar in each experimental condition; these lysates were then incubated at 37°C overnight. To reduce non-specific binding, lysates were pre-cleared with streptavidin beads and then were incubated for 2 h at 30°C with 4 μg biotinylated *E. coli* O111:B4 LPS loaded onto 8 μg of Streptavidin Mag sepharose (GE, 28985738, or Pierce, 88816).

### Plate binding assays

Clear flat bottom Nunc Maxisorp ELISA plates (Thermo Fisher, 442404) were coated with various concentrations of purified recombinant caspase-4 (Origene, TP760359) overnight at 4°C. The plates were washed 3x with PBS then blocked for 2 h at RT with 1% bovine serum albumin (Fisher, BP9700100) in PBS. Excess blocking solution was removed. HEK 293T cells transiently expressing FLAG-NLRP11 and control untransfected cells were lysed into 50 mM Tris-HCl pH 7.4, 150 mM NaCl, 5 mM MgCl_2_, 100 µg/ml digitonin (Thermo Fisher Scientific, BN2006) and complete Mini EDTA-free protease inhibitor (Roche), as described (*30*). Total protein concentration of lysates was normalized to 2 mg/mL. Serial dilutions were in untransfected HEK293T cell lysate. Varying concentrations of cell lysates were incubated with pre-blocked ELISA plates for 1 h at RT. Excess lysate was removed by washing 3 times with PBS. For detection of FLAG-NLRP11, wells were incubated with 0.8 μg/mL anti-FLAG rabbit (Sigma, F7425-.2MG) was incubated for 1 h at RT, and after washing with PBS, 0.16 μg/mL goat-anti-rabbit conjugated to horseradish peroxidase (HRP) (Jackson, 111-035-144). To detect caspase-4 bound to the plate, wells were incubated with 0.5 μg/mL anti-caspase-4 (Santa Cruz, sc-56056) and, after washing with PBS, 0.16 μg/mL goat-anti-mouse conjugated to HRP (Jackson, 115-035-003), each for 1 hr. After 6 washes with PBS, 50 μL of One Step Ultra-TMB (Fisher Scientific, PI34029) was added to each well and incubated for 5 min before stopping with 4 N sulfuric acid and reading in a BioTek plate reader.

### Bacterial infection

Overnight cultures were started from a single colony taken from a freshly streaked plate. Infections with *S. flexneri* were performed with exponential phase bacteria, after back-dilution from overnight cultures. Infections with *L. monocytogenes* were performed with stationary phase bacteria from overnight cultures. Infections were with a multiplicity of infection (MOI) of 10 for *S. flexneri* cultures and 1 for *L. monocytogenes*. For *L. monocytogenes* infections, bacteria were mixed with 10 ug/ml LPS or no LPS in a volume of 1 ml just prior to infection of the cells (*13*). Bacteria were centrifuged onto cells at 2000 rpm for 10 min, followed by incubation at 37°C. After 30 min, the cells were washed twice, and then 25 μg/mL gentamicin was added to the media to kill extracellular bacteria. To determine efficiency of bacterial invasion, cells were harvested with trypsin 20 minutes later and lysed with 0.5% Triton X-100, and lysates were plated for bacterial counts. Where appropriate *S. enterica* serovar Minnesota LPS was added to 10 μg/mL at the time of infection and MCC950 to 1 μM 30 min prior to infection and maintained during infection. For LDH assays of *S. flexneri* infected cells, 100 μL of cell culture supernatant was collected at 2 h and 30 min and at 5 h and/or 30 min of infection; for detection of caspases and/or gasdermin D by western blot, 100 μL of cell culture supernatant was collected at the same time points and was mixed with 35 μL 4x Laemmli buffer supplemented with cOmplete EDTA-free protease inhibitors (Roche, 5056489001). For parallel analyses of *L. monocytogenes* infected cells, gentamicin was used at 15 μg/mL, and samples were collected at 6 h of infection. For western blot analyses, whole cell lysates were collected in 2x Laemmli buffer.

Infections with *B. thailandensis* and *S*. Typhimurium were performed with exponential phase bacteria at a multiplicity of infection (MOI) of 10. Bacteria were centrifuged onto cells at 2000 rpm for 10 min, followed by incubation at 37°C. For *S*. Typhimurium, after 30 min incubation, the cells were washed twice, and then 25 μg/mL gentamicin was added to the media to kill extracellular bacteria. For *B. thailandensis*, after 2 h incubation, the cells were washed twice with cRPMI. For LDH assays with *S*. Typhimurium infection, 100 μL of cell culture supernatant was collected at 5 h 30 min to 7 h 30 min of infection and, for LDH assays with *B. thailandensis*, 100 μL of cell culture supernatant was collected at 24 h.

### Isolation of primary human peripheral blood mononuclear cells (PBMCs) and CD14 monocytes

Cells were isolated from leukopaks from individual healthy donors (Crimson Core Laboratories, Brigham and Women’s Hospital) using density-gradient centrifugation. Briefly, blood was diluted 1:2 with RPMI-1640 (Gibco), layered on top of Ficoll-Paque Plus (GE Healthcare) using SepMate™-50 (IVD) (StemCell Technologies), and centrifuged at 1,200g for 20 min. The PBMC layer was collected and resuspended in 50 ml RPMI-1640 (Gibco) and centrifuged again at 300 g for 10 min. The cells were counted, resuspended in Cryostor CS10 (StemCell Technologies), and aliquoted in cryopreservation tubes at a concentration of 1 × 10^8^ cells per mL. The tubes were placed at −80°C overnight, then transferred to liquid nitrogen for long-term storage. To isolate CD14+ cells, EasySep™ Human CD14 Positive Selection Kit II (StemCell Technologies) was used, following the manufacturer’s instructions. Cells were cultured in complete RPMI, differentiated with 25 ng/mL m-CSF (StemCell Technologies), and maintained at 37°C in 5% CO_2_.

### LPS electroporation

Electroporation of cells with LPS was performed, as previously described (*14*), using the Neon Transfection System (Invitrogen) according to the manufacturer’s instructions using *S*. Minnesota LPS at indicated concentrations. THP-1 cells were seeded in low-attachment 10 cm dishes with 50 ng/mL PMA in RPMI. After 48 h, the cells were lifted in PBS (without calcium or magnesium). For electroporation, 1.5 × 10^6^ THP-1 cells were pelleted, resuspended in Resuspension Buffer R with 2 μg/mL of LPS or water control. Using a 10 μL Neon tip, cells were pulsed once at 1400 V and 10 ms pulse width. For primary human macrophage cells, 7.5 × 10^5^ CD14 macrophages were pelleted, then resuspended in Resuspension Buffer T with 2 μg/mL of LPS or water control. Using a 10 μL Neon tip, cells were pulsed 2 times at 2250 V and 20 ms pulse width. Electroporated cells were immediately resuspended in 2 mL of RPMI and placed into wells of a 24 well plate. Cells were incubated for 2.5 h at 37°C before measuring LDH release.

### siRNA transient knockdowns

siRNAs used for NLRP11 knockdowns were Hs_NALP11_1 FlexiTube siRNA and Hs_NALP11_5 FlexiTube siRNA at 10 nM total (5 nM each siRNA); for caspase-4 knockdowns were 3941, 4036 and 4130 at 30 nM total (10 nM each siRNA) (ThermoFisher); and for Caspase-1 Hs_CASP1_14 FlexiTube siRNA and Hs_CASP1_13 FlexiTube siRNA at 30 nM total (15 nM each siRNA). AllStars Negative Control siRNAs (Qiagen) were used as negative controls at 10 nM or 30 nM, accordingly. In each set of experiments, to control for loading, an identical number of cells were used and processed for each experimental condition. siRNAs were transfected using the HiPerfect reagent (Qiagen) into primary human monocyte-derived macrophages 72 h before or into THP-1 cells 48 h before experimental assays, following the manufacturer’s protocol for “Transfection of Differentiated Macrophage Cell Lines.” After transfection, cells were maintained at 37°C in 5% CO_2_. Degree of knock-down of caspases was assessed by western blot of lysates of equivalent numbers of cells for each condition, and of *NLRP11* by reverse transcriptase (RT)-PCR. For RT-PCR, cellular transcripts were prepared using RNeasy Micro Kit (Qiagen) on cells pretreated or not with Pam3CSK4, cDNA was generated using SuperScript IV VILO kit (ThermoFisher), and PCR was performed for 40 cycles.

### LDH assay and cytokine ELISAs

LDH was measured using an LDH Cytotoxicity Kit (Pierce, 88953) or Cytotoxicity Detection Kit (Sigma, 11644793001) per the manufacturer’s instructions; for any set of experiments, the same assay kit was used for all experiments. Measurements at 490 nm and 680 nm were taken immediately after addition of Stop Solution using a BioTek Plate Reader. To determine LDH activity, the 680 nm absorbance value (background) was subtracted from the 490-nm absorbance value, and cytotoxicity was calculated for each sample as LDH activity as a fraction of the total LDH activity for cells treated with the Triton X-100 lysis buffer provided by the manufacturer. For experiments involving primary cells, a different donor was used for each biological replicate; due to inherent variability from donor to donor in the baseline levels of death in cell death assays, within-donor data were normalized to the negative control condition before compilation of the biological replicates. Assays of levels of secreted cytokines were performed using the Human Total IL-18 DuoSet ELISA (R&D Biosystems, DY318-05), ELISA MAX Deluxe Set Human IL-1a (Biolegend, 445804), and ELISA MAX Deluxe Set Human IL-1b (Biolegend, 437004), per the manufacturer’s instructions. Where applicable, as a positive control, cell culture supernatants were also analyzed following priming with 1 μM O55:B5 *E. coli* LPS for 3 h, at which time 10 μM nigericin was added; LDH release was assayed 2 h after addition of nigericin.

### Statistics and reproducibility

Unless otherwise indicated, all results are data from at least 3 independent biological replicates. On bar graphs, each individual symbol is data from an independent biological replicate. Statistical analysis was performed with Prism version 9.

## Data availability

All relevant data are included in the manuscript and will be made available by request to the corresponding author.

## List of Supplementary Materials

Figs. S1 to S12

## Acknowledgments

We thank Jonathan Kagan, Hao Wu, Kate Fitzgerald, Sunny Shin, Ann Goldfeld, Margaret Kielian, Peggy Cotter, and Jeffrey Miller for reagents.

## Funding

Real Colegio Complutense Fellowship, Real Colegio Complutense at Harvard (MLGM)

National Institutes of Health grant R01 AI081724 (MBG)

National Institutes of Health grant F32 AI145128 (ASZ)

National Institutes of Health grant F32 AI126765 (KAM)

National Institutes of Health grant T32 AI007061 (ASZ)

## Author contributions

Conceptualization: MBG

Methodology: MRL, ASZ, TEW, MBG

Investigation: MRL, ASZ, TEW, KAM, MLGM, ACH, VK, KTE

Funding acquisition: ASZ, KAM, MLGM, MBG

Supervision: MRL, KAM, MBG

Writing – original draft: MBG

Writing – review & editing: MRL, ASZ, TEW, KAM, MLGM, MBG

## Competing interests

The authors declare no competing interests.

## Data and materials availability

All data are available in the main text or the supplementary materials. All materials are available by request to MBG, restricted by institutional MTAs.

## Notes

### Competing Interest Statement

The authors have declared no competing interest.

## References and Notes

1. S. M. Brewer, S. W. Brubaker, D. M. Monack, Host inflammasome defense mechanisms and bacterial pathogen evasion strategies. Curr Opin Immunol 60, 63–70 (2019).

2. P. Broz, V. M. Dixit, Inflammasomes: mechanism of assembly, regulation and signalling. Nat Rev Immunol 16, 407–420 (2016).

3. I. Jorgensen, M. Rayamajhi, E. A. Miao, Programmed cell death as a defence against infection. Nat Rev Immunol 17, 151–164 (2017).

4. S. S. Wright, S. O. Vasudevan, V. A. Rathinam, Mechanisms and Consequences of Noncanonical Inflammasome-Mediated Pyroptosis. J Mol Biol, 167245 (2021).

5. K. A. Deets, R. E. Vance, Inflammasomes and adaptive immune responses. Nat Immunol 22, 412–422 (2021).

6. N. Kayagaki, I. B. Stowe, B. L. Lee, K. O’Rourke, K. Anderson, S. Warming, T. Cuellar, B. Haley, M. Roose-Girma, Q. T. Phung, P. S. Liu, J. R. Lill, H. Li, J. Wu, S. Kummerfeld, J. Zhang, W. P. Lee, S. J. Snipas, G. S. Salvesen, L. X. Morris, L. Fitzgerald, Y. Zhang, E. M. Bertram, C. C. Goodnow, V. M. Dixit, Caspase-11 cleaves gasdermin D for non-canonical inflammasome signalling. Nature 526, 666–671 (2015).

7. J. Shi, Y. Zhao, K. Wang, X. Shi, Y. Wang, H. Huang, Y. Zhuang, T. Cai, F. Wang, F. Shao, Cleavage of GSDMD by inflammatory caspases determines pyroptotic cell death. Nature 526, 660–665 (2015).

8. J. Ding, K. Wang, W. Liu, Y. She, Q. Sun, J. Shi, H. Sun, D. C. Wang, F. Shao, Pore-forming activity and structural autoinhibition of the gasdermin family. Nature 535, 111–116 (2016).

9. W. T. He, H. Wan, L. Hu, P. Chen, X. Wang, Z. Huang, Z. H. Yang, C. Q. Zhong, J. Han, Gasdermin D is an executor of pyroptosis and required for interleukin-1beta secretion. Cell Res 25, 1285–1298 (2015).

10. X. Liu, Z. Zhang, J. Ruan, Y. Pan, V. G. Magupalli, H. Wu, J. Lieberman, Inflammasome-activated gasdermin D causes pyroptosis by forming membrane pores. Nature 535, 153–158 (2016).

11. N. Kayagaki, S. Warming, M. Lamkanfi, L. Vande Walle, S. Louie, J. Dong, K. Newton, Y. Qu, J. Liu, S. Heldens, J. Zhang, W. P. Lee, M. Roose-Girma, V. M. Dixit, Non-canonical inflammasome activation targets caspase-11. Nature 479, 117–121 (2011).

12. N. Kayagaki, M. T. Wong, I. B. Stowe, S. R. Ramani, L. C. Gonzalez, S. Akashi-Takamura, K. Miyake, J. Zhang, W. P. Lee, A. Muszynski, L. S. Forsberg, R. W. Carlson, V. M. Dixit, Noncanonical inflammasome activation by intracellular LPS independent of TLR4. Science 341, 1246–1249 (2013).

13. J. A. Hagar, D. A. Powell, Y. Aachoui, R. K. Ernst, E. A. Miao, Cytoplasmic LPS activates caspase-11: implications in TLR4-independent endotoxic shock. Science 341, 1250–1253 (2013).

14. J. Shi, Y. Zhao, Y. Wang, W. Gao, J. Ding, P. Li, L. Hu, F. Shao, Inflammatory caspases are innate immune receptors for intracellular LPS. Nature 514, 187–192 (2014).

15. B. C. Russo, L. M. Stamm, M. Raaben, C. M. Kim, E. Kahoud, L. R. Robinson, S. Bose, A. L. Queiroz, B. B. Herrera, L. A. Baxt, N. Mor-Vaknin, Y. Fu, G. Molina, D. M. Markovitz, S. P. Whelan, M. B. Goldberg, Intermediate filaments enable pathogen docking to trigger type 3 effector translocation. Nat Microbiol 1, 16025 (2016).

16. K. Ellwanger, E. Becker, I. Kienes, A. Sowa, Y. Postma, Y. Cardona Gloria, A. N. R. Weber, T. A. Kufer, The NLR family pyrin domain-containing 11 protein contributes to the regulation of inflammatory signaling. The Journal of biological chemistry 293, 2701–2710 (2018).

17. C. Wu, Z. Su, M. Lin, J. Ou, W. Zhao, J. Cui, R. F. Wang, NLRP11 attenuates Toll-like receptor signalling by targeting TRAF6 for degradation via the ubiquitin ligase RNF19A. Nat Commun 8, 1977 (2017).

18. Y. Qin, Z. Su, Y. Wu, C. Wu, S. Jin, W. Xie, W. Jiang, R. Zhou, J. Cui, NLRP11 disrupts MAVS signalosome to inhibit type I interferon signaling and virus-induced apoptosis. EMBO Rep 18, 2160–2171 (2017).

19. T. Sanada, M. Kim, H. Mimuro, M. Suzuki, M. Ogawa, A. Oyama, H. Ashida, T. Kobayashi, T. Koyama, S. Nagai, Y. Shibata, J. Gohda, J. Inoue, T. Mizushima, C. Sasakawa, The Shigella flexneri effector OspI deamidates UBC13 to dampen the inflammatory response. Nature 483, 623–626 (2012).

20. S. Ruhl, P. Broz, Caspase-11 activates a canonical NLRP3 inflammasome by promoting K(+) efflux. Eur J Immunol 45, 2927–2936 (2015).

21. J. L. Schmid-Burgk, M. M. Gaidt, T. Schmidt, T. S. Ebert, E. Bartok, V. Hornung, Caspase-4 mediates non-canonical activation of the NLRP3 inflammasome in human myeloid cells. Eur J Immunol 45, 2911–2917 (2015).

22. V. A. Rathinam, S. K. Vanaja, L. Waggoner, A. Sokolovska, C. Becker, L. M. Stuart, J. M. Leong, K. A. Fitzgerald, TRIF licenses caspase-11-dependent NLRP3 inflammasome activation by gram-negative bacteria. Cell 150, 606–619 (2012).

23. B. A. Napier, S. W. Brubaker, T. E. Sweeney, P. Monette, G. H. Rothmeier, N. A. Gertsvolf, A. Puschnik, J. E. Carette, P. Khatri, D. M. Monack, Complement pathway amplifies caspase-11-dependent cell death and endotoxin-induced sepsis severity. J Exp Med 213, 2365–2382 (2016).

24. P. J. Baker, D. Boucher, D. Bierschenk, C. Tebartz, P. G. Whitney, D. B. D’Silva, M. C. Tanzer, M. Monteleone, A. A. Robertson, M. A. Cooper, S. Alvarez-Diaz, M. J. Herold, S. Bedoui, K. Schroder, S. L. Masters, NLRP3 inflammasome activation downstream of cytoplasmic LPS recognition by both caspase-4 and caspase-5. Eur J Immunol 45, 2918–2926 (2015).

25. N. J. Bitto, P. J. Baker, J. K. Dowling, G. Wray-McCann, A. De Paoli, L. S. Tran, P. L. Leung, K. J. Stacey, A. Mansell, S. L. Masters, R. L. Ferrero, Membrane vesicles from Pseudomonas aeruginosa activate the noncanonical inflammasome through caspase-5 in human monocytes. Immunol Cell Biol 96, 1120–1130 (2018).

26. A. Gangopadhyay, S. Devi, S. Tenguria, J. Carriere, H. Nguyen, E. Jager, H. Khatri, L. H. Chu, R. A. Ratsimandresy, A. Dorfleutner, C. Stehlik, NLRP3 licenses NLRP11 for inflammasome activation in human macrophages. Nat Immunol 23, 892–903 (2022).

27. D. M. Pilla, J. A. Hagar, A. K. Haldar, A. K. Mason, D. Degrandi, K. Pfeffer, R. K. Ernst, M. Yamamoto, E. A. Miao, J. Coers, Guanylate binding proteins promote caspase-11-dependent pyroptosis in response to cytoplasmic LPS. Proc Natl Acad Sci U S A 111, 6046–6051 (2014).

28. D. Fisch, H. Bando, B. Clough, V. Hornung, M. Yamamoto, A. R. Shenoy, E. M. Frickel, Human GBP1 is a microbe-specific gatekeeper of macrophage apoptosis and pyroptosis. The EMBO journal 38, e100926 (2019).

29. J. C. Santos, M. S. Dick, B. Lagrange, D. Degrandi, K. Pfeffer, M. Yamamoto, E. Meunier, P. Pelczar, T. Henry, P. Broz, LPS targets host guanylate-binding proteins to the bacterial outer membrane for non-canonical inflammasome activation. The EMBO journal 37, (2018).

30. M. P. Wandel, B. H. Kim, E. S. Park, K. B. Boyle, K. Nayak, B. Lagrange, A. Herod, T. Henry, M. Zilbauer, J. Rohde, J. D. MacMicking, F. Randow, Guanylate-binding proteins convert cytosolic bacteria into caspase-4 signaling platforms. Nat Immunol 21, 880–891 (2020).

31. J. C. Santos, D. Boucher, L. K. Schneider, B. Demarco, M. Dilucca, K. Shkarina, R. Heilig, K. W. Chen, R. Y. H. Lim, P. Broz, Human GBP1 binds LPS to initiate assembly of a caspase-4 activating platform on cytosolic bacteria. Nat Commun 11, 3276 (2020).

32. B. H. Kim, A. R. Shenoy, P. Kumar, C. J. Bradfield, J. D. MacMicking, IFN-inducible GTPases in host cell defense. Cell host & microbe 12, 432–444 (2012).

33. M. Kutsch, L. Sistemich, C. F. Lesser, M. B. Goldberg, C. Herrmann, J. Coers, Direct binding of polymeric GBP1 to LPS disrupts bacterial cell envelope functions. The EMBO journal 39, e104926 (2020).

34. J. L. Poyet, S. M. Srinivasula, M. Tnani, M. Razmara, T. Fernandes-Alnemri, E. S. Alnemri, Identification of Ipaf, a human caspase-1-activating protein related to Apaf-1. The Journal of biological chemistry 276, 28309–28313 (2001).

35. C. Pop, J. Timmer, S. Sperandio, G. S. Salvesen, The apoptosome activates caspase-9 by dimerization. Mol Cell 22, 269–275 (2006).

36. R. A. MacCorkle, K. W. Freeman, D. M. Spencer, Synthetic activation of caspases: artificial death switches. Proc Natl Acad Sci U S A 95, 3655–3660 (1998).

37. B. Faustin, L. Lartigue, J. M. Bruey, F. Luciano, E. Sergienko, B. Bailly-Maitre, N. Volkmann, D. Hanein Rouiller, J. C. Reed, Reconstituted NALP1 inflammasome reveals two-step mechanism of caspase-1 activation. Mol Cell 25, 713–724 (2007).

38. B. C. Baliga, S. H. Read, S. Kumar, The biochemical mechanism of caspase-2 activation. Cell Death Differ 11, 1234–1241 (2004).

39. P. Karki, G. R. Dahal, I. S. Park, Both dimerization and interdomain processing are essential for caspase-4 activation. Biochemical and biophysical research communications 356, 1056–1061 (2007).

40. K. Wang, Q. Sun, X. Zhong, M. Zeng, H. Zeng, X. Shi, Z. Li, Y. Wang, Q. Zhao, F. Shao, J. Ding, Structural Mechanism for GSDMD Targeting by Autoprocessed Caspases in Pyroptosis. Cell 180, 941–955 e920 (2020).

41. Q. Yin, H. H. Park, J. Y. Chung, S. C. Lin, Y. C. Lo, L. S. da Graca, X. Jiang, H. Wu, Caspase-9 holoenzyme is a specific and optimal procaspase-3 processing machine. Mol Cell 22, 259–268 (2006).

42. H. Sasaki, S. H. White, Aggregation behavior of an ultra-pure lipopolysaccharide that stimulates TLR-4 receptors. Biophys J 95, 986–993 (2008).

43. M. A. Wacker, A. Teghanemt, J. P. Weiss, J. H. Barker, High-affinity caspase-4 binding to LPS presented as high molecular mass aggregates or in outer membrane vesicles. Innate Immun 23, 336–344 (2017).

44. A. M. Taveira da Silva, H. C. Kaulbach, F. S. Chuidian, D. R. Lambert, A. F. Suffredini, R. L. Danner, Brief report: shock and multiple-organ dysfunction after self-administration of Salmonella endotoxin. N Engl J Med 328, 1457–1460 (1993).

45. R. W. Schaedler, R. J. Dubos, The susceptibility of mice to bacterial endotoxins. J Exp Med 113, 559–570 (1961).

46. E. H. Labrec, H. Schneider, T. J. Magnani, S. B. Formal, Epithelial Cell Penetration as an Essential Step in the Pathogenesis of Bacillary Dysentery. J Bacteriol 88, 1503–1518 (1964).

47. H. J. Wing, A. W. Yan, S. R. Goldman, M. B. Goldberg, Regulation of IcsP, the outer membrane protease of the Shigella actin tail assembly protein IcsA, by virulence plasmid regulators VirF and VirB. J Bacteriol 186, 699–705 (2004).

48. A. T. Maurelli, B. Blackmon, R. Curtiss, 3rd, Loss of pigmentation in Shigella flexneri 2a is correlated with loss of virulence and virulence-associated plasmid. Infect Immun 43, 397–401 (1984).

49. K. A. Miller, A. C. Garza-Mayers, Y. Leung, M. B. Goldberg, Identification of interactions among host and bacterial proteins and evaluation of their role early during Shigella flexneri infection. Microbiology (Reading) 164, 540–550 (2018).

50. C. R. Beuzon, S. Meresse, K. E. Unsworth, J. Ruiz-Albert, S. Garvis, S. R. Waterman, T. A. Ryder, E. Boucrot, D. W. Holden, Salmonella maintains the integrity of its intracellular vacuole through the action of SifA. The EMBO journal 19, 3235–3249 (2000).

51. V. Novem, G. Shui, D. Wang, A. K. Bendt, S. H. Sim, Y. Liu, T. W. Thong, S. P. Sivalingam, E. E. Ooi, M. R. Wenk, G. Tan, Structural and biological diversity of lipopolysaccharides from Burkholderia pseudomallei and Burkholderia thailandensis. Clin Vaccine Immunol 16, 1420–1428 (2009).

52. M. M. Gedde, D. E. Higgins, L. G. Tilney, D. A. Portnoy, Role of listeriolysin O in cell-to-cell spread of Listeria monocytogenes. Infect Immun 68, 999–1003 (2000).

53. K. A. Datsenko, B. L. Wanner, One-step inactivation of chromosomal genes in Escherichia coli K-12 using PCR products. Proc Natl Acad Sci U S A 97, 6640–6645 (2000).

54. E. K. Brinkman, T. Chen, M. Amendola, B. van Steensel, Easy quantitative assessment of genome editing by sequence trace decomposition. Nucleic Acids Res 42, e168 (2014).

55. S. Ranjbar, L. D. Jasenosky, N. Chow, A. E. Goldfeld, Regulation of Mycobacterium tuberculosis-dependent HIV-1 transcription reveals a new role for NFAT5 in the toll-like receptor pathway. PLoS Pathog 8, e1002620 (2012).

56. T. Kobayashi, M. Ogawa, T. Sanada, H. Mimuro, M. Kim, H. Ashida, R. Akakura, M. Yoshida, M. Kawalec, J. M. Reichhart, T. Mizushima, C. Sasakawa, The Shigella OspC3 effector inhibits caspase-4, antagonizes inflammatory cell death, and promotes epithelial infection. Cell host & microbe 13, 570–583 (2013).

57. Z. Li, W. Liu, J. Fu, S. Cheng, Y. Xu, Z. Wang, X. Liu, X. Shi, Y. Liu, X. Qi, X. Liu, J. Ding, F. Shao, Shigella evades pyroptosis by arginine ADP-riboxanation of caspase-11. Nature 599, 290–295 (2021).

