## Supplementary materials for "Pattern Recognition Receptor for Bacterial Lipopolysaccharide in the Cytosol of Human Macrophages"

**This pdf file includes:**

Figs. S1 to S12

### Supplementary Figures

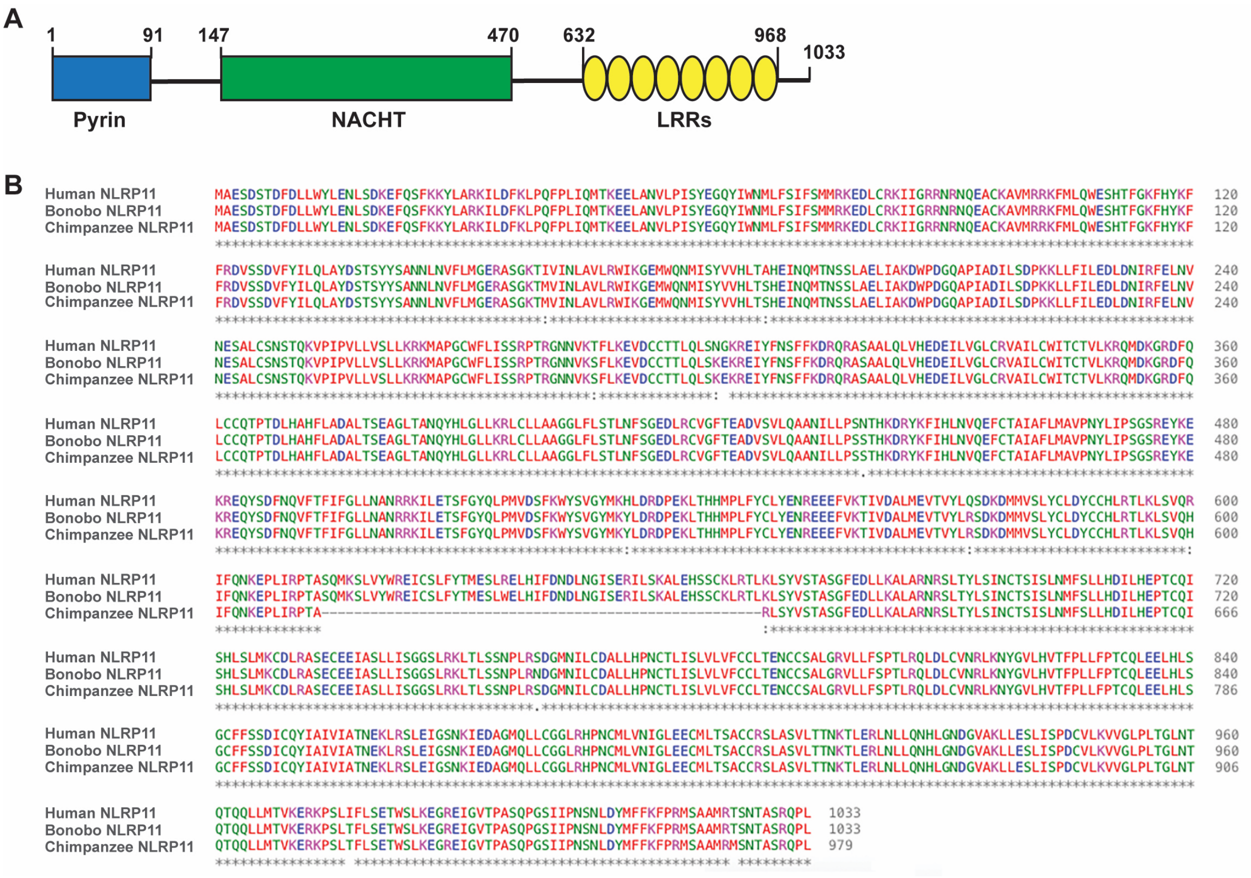


**Fig. S1. Human NLRP11 and primate homologs.**

(**A**) Domain organization of NLRP11; proposed amino acid delimitations (above solid shapes) are based on alignment with NALP4, the human NLRP to which NLRP11 is most similar (E-value 0.0; score 1511; 35.0% identical and 54.4% conserved residues over 994 match length). LRR, leucine-rich repeat. (**B**) Alignment of human NLRP11 (NAL11_HUMAN) isoform 1 with Bonobo NLRP11 (A0A2R8ZGR4_PANPA) and chimpanzee NLRP11 (A0A2I3RD68_PANTR); sequences from UniProt.

A

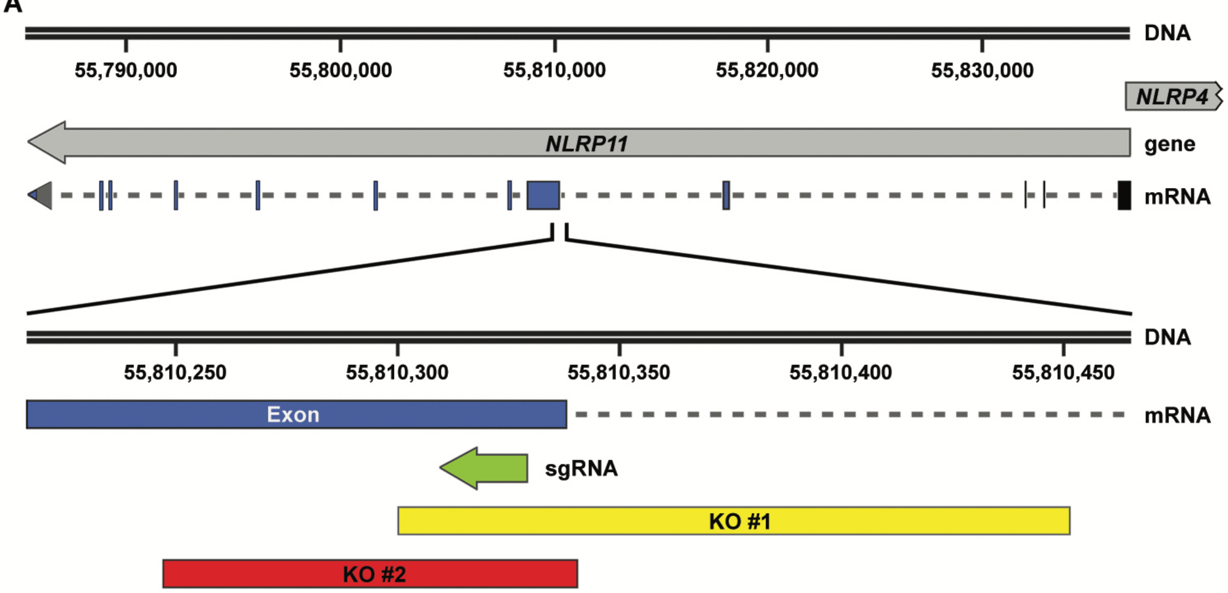


B

| | Coordinates | Gene ID (NCBI) | Location |
|---|---|---|---|
| *NLRP11* | 55,785,397..55,836,762, complement | 204801 | 19q13.42-q13.43 |
| *NLRP4* | 55,836,578..55,881,855 | 147945 | 19q13.43 |
| *NLRP11* sgRNA | 55,810,310..55,810,329, complement | 204801 | |
| *NLRP11* KO #1 | Deletion 55,810,301..55,810,451 | 204801 | |
| *NLRP11* KO #2 | Deletion 55,810,248..55,810,340 | 204801 | |

C

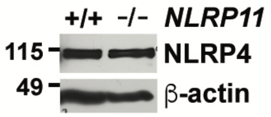


**Fig. S2. Generation and validation of *NLRP11* deletions.**

(**A**) Map of human genomic context of *NLRP11* gene, on the complementary strand, and *NLRP4*, which overlaps by 185 nucleotides and is oriented in the opposite direction on the sense strand. Also shown are *NLRP11* protein coding exons (blue bars) and non-coding exons (black bars), the guide RNA used to generate *NLRP11* deletions (sgRNA, green arrow), and the extent of the deletions in the two THP-1 *NLRP11* deletion cell lines used in this study (KO #1 and KO #2, yellow and red bars, respectively). Double lines represent DNA and dashed line represents intron mRNA. Numbering under the double lines indicates genome coordinates in NCBI Reference Sequence NC_000019.10. (**B**) Coordinates of relevant genes and deletions. Each *NLRP11*$^{-/-}$ knock-out (KO) clone is homozygous for the indicated deletion. (**C**) Production of NLRP4 is unaltered by the deletion in *NLRP11*. Shown are cell lysates of THP-1 parent cell line and THP-1 *NLRP11* KO #1. Western blot for NLRP4; *n* = 1 experiment; β-actin, loading control. MW markers in kD.

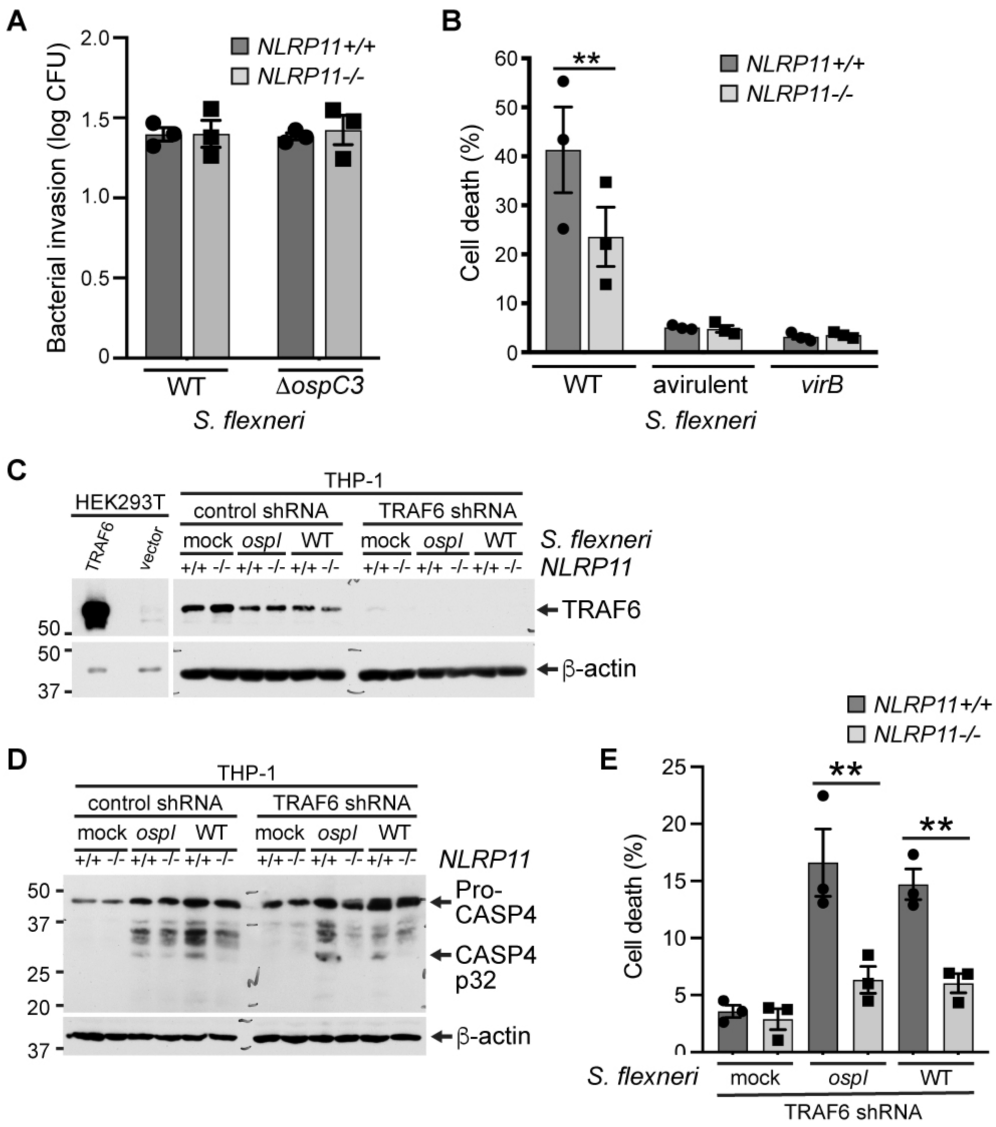


**Fig. S3. *S. flexneri* invasion, the type 3 secretion system, and TRAF6 in bacterial infection-induced death of THP-1 macrophage-like cells lacking NLRP11.**

(**A**) Efficiency of invasion of indicated *S. flexneri* strains into *NLRP11*$^{+/+}$ or *NLRP11*$^{-/-}$ monocyte-derived THP-1 macrophages. (**B**) *S. flexneri* that lack type 3 secretion function by virtue of lacking the virulence plasmid (avirulent) or carrying a deletion of the virulence regulator *virB* induce negligible cell death in human-derived THP-1 macrophage-like cells, whether or not NLRP11 is present. *n* = 3 experiments. **, $p<0.05$. Two-way ANOVA with Sidak *post hoc* test. (**C-E**) *S. flexneri*-induced NLRP11-dependent cell death is independent of TRAF6. Caspase-4 (CASP4) activation and cell death following *S. flexneri* infection of *NLRP11*$^{-/-}$ or *NLRP11*$^{+/+}$ monocyte-derived THP-1 macrophages, stably transfected or not with TRAF6 shRNA. *S. flexneri* wildtype (WT) or *ospI* mutant, or mock-infected. TRAF6 levels, with β-actin loading control (**C**), CASP4 processing (**D**), and cell death, measured by LDH release (**E**). In (**C**), the two TRAF6 panels are from the same blot, and the two β-actin panels are from the same blot. Representative western blots of cell lysates (both blots in panel **C** and β-actin blot in panel **D**) or culture supernatants (CASP4 blot in panel **D**). Cell death measured as LDH release relative to complete lysis control. *n* = 3 experiments. MW markers in kD. Two-way ANOVA with Sidak *post hoc* test (**B**) or Tukey *post hoc* test (**E**). Mean ± S.E.M. **, $p<0.01$.

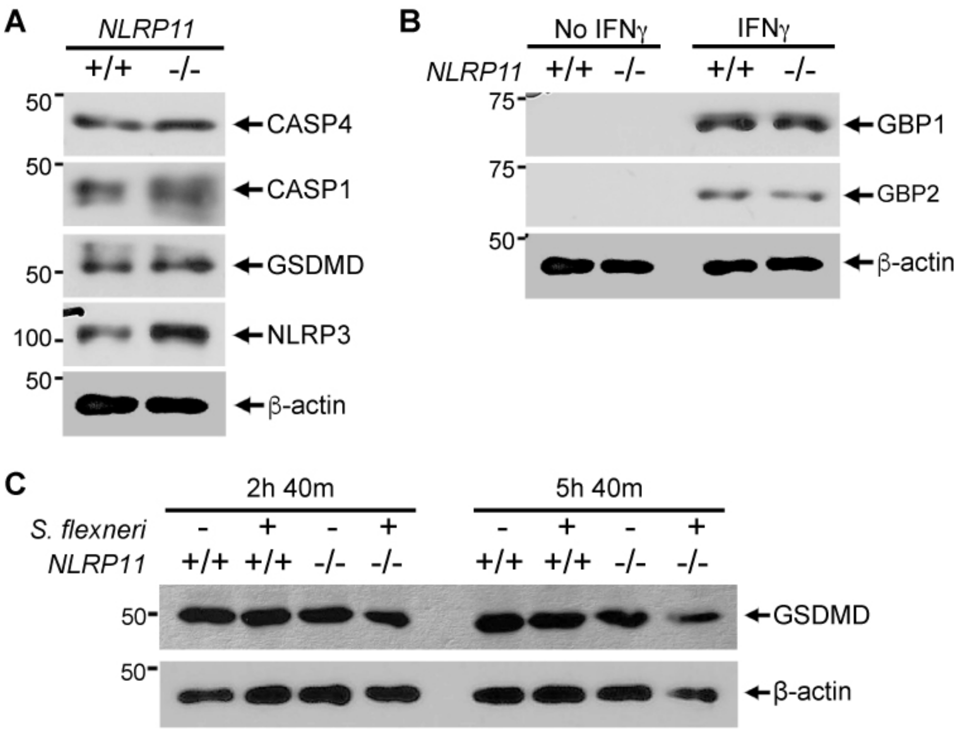


**Fig. S4. Endogenous levels of innate immune proteins in *NLRP11*$^{-/-}$ macrophage-like THP-1 cells.**

(**A**) Endogenous levels of indicated innate immune proteins in *NLRP11*$^{+/+}$ and *NLRP11*$^{-/-}$ macrophage-line THP-1 cells. Representative western blots of cell lysates. *n* = 2 experiments. (**B**) Endogenous levels of guanylate binding proteins 1 and 2 in *NLRP11*$^{+/+}$ and *NLRP11*$^{-/-}$ macrophage-line THP-1 cells with or without pre-treatment with interferon (IFN)γ. Representative western blots of cell lysates. *n* = >3 experiments. (**C**) Levels of gasdermin D in *S. flexneri*-infected or uninfected *NLRP11*$^{+/+}$ and *NLRP11*$^{-/-}$ macrophage-line THP-1 cells, at indicated durations of infection. *n* = >3 experiments. MW markers in kD. CASP4, caspase-4; CASP1, caspase-1; GSDMD, gasdermin D; GBP1, guanylate binding protein 1; GBP2, guanylate binding protein 2.

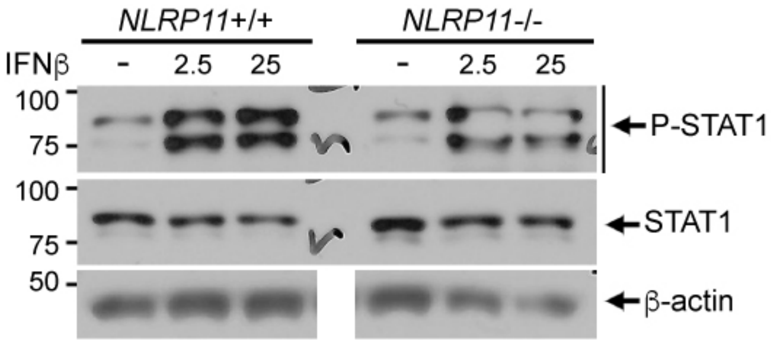


**Fig. S5. Response of NLRP11-/- cells to type I interferon stimulation.**

Levels of phosphorylation of STAT1 in *NLRP11*$^{+/+}$ and *NLRP11*$^{-/-}$ macrophage-like THP-1 cells following pre-treatment or no pre-treatment with interferon (IFN)-β at concentration (in U/mL) indicated above blots. Representative western blots. β-actin, loading control. MW markers in kD. *n*=3 experiments.

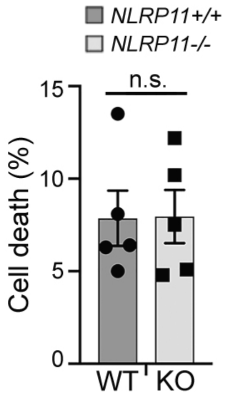


**Fig. S6. Cell death associated with *NLRP11* is independent of extracellular LPS.**

Cell death of *NLRP11*$^{-/-}$ and *NLRP11*$^{+/+}$ THP-1 macrophage-like cells upon exposure to extracellular LPS (2 µg/ml) for 24 h. Two-way ANOVA, p = 0.99. *n* = 5 experiments. Paired student's two-tailed T-test. Mean ± S.E.M. n.s., not significant.

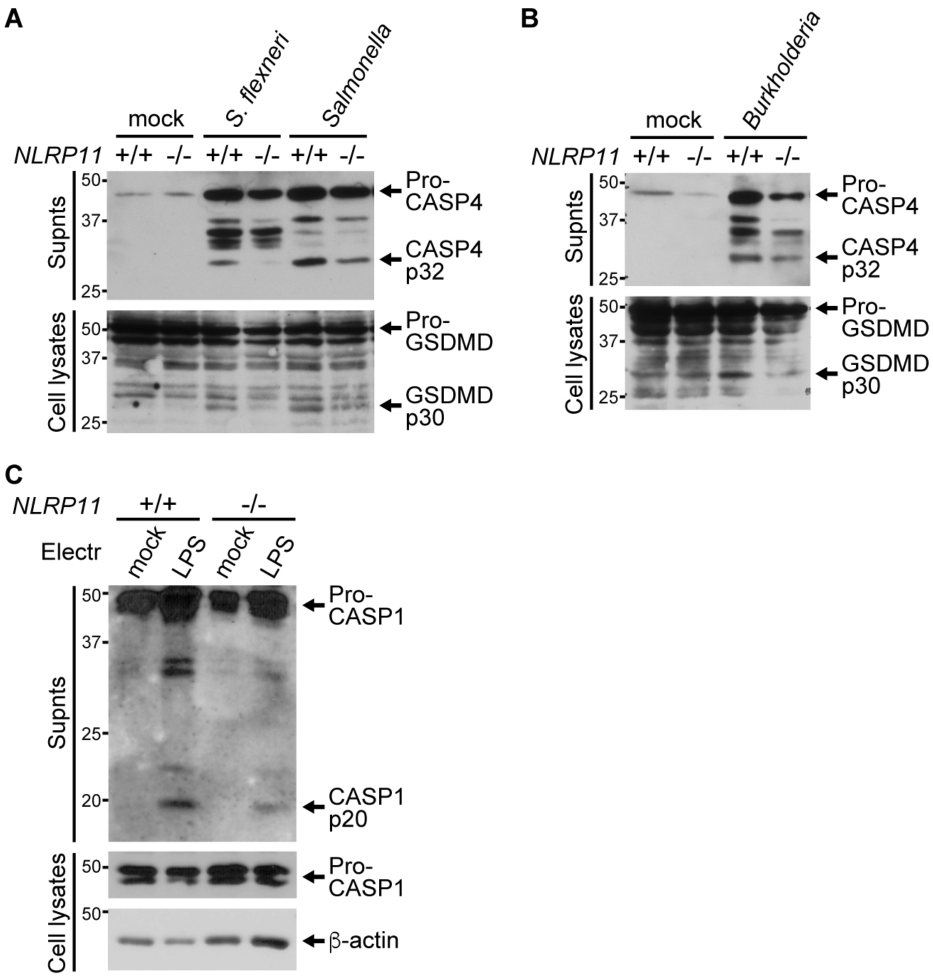


**Fig. S7. Processing of caspases upon bacterial infection in *NLRP11*$^{-/-}$ THP-1 macrophage-like cells.**

(**A-B**) Processing of caspase-4 (CASP4), with release of p32 polypeptide (top), and gasdermin D (GSDMD), with liberation of p30 polypeptide (bottom) upon infection of THP-1 macrophages with *S. flexneri* or *S.* Typhimurium (**A**) or with *B. thailandensis* (**B**). Representative western blots of culture supernatants (Supnts, CASP4) or associated cell lysates (GSDMD). (**C**) Processing of caspase-1 (CASP1), with release of p20 polypeptide, upon electroporation (Electr) of THP-1 macrophages with LPS. Representative western blots of released caspase-1 in culture supernatants (CASP1) and caspase-1 and β-actin control in associated cell lysates. MW markers in kD.

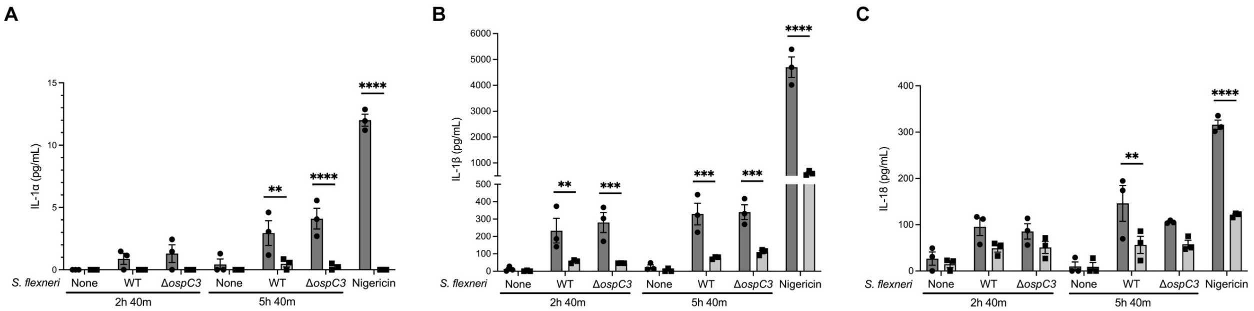


**Fig. S8. NLRP11-dependence of release of IL-1 cytokines from THP-1 macrophage-like cells upon *S. flexneri* infection.**

Release of IL-1α, Il-1β, and IL-18 from WT and *NLRP11*$^{-/-}$ macrophage-like THP-1 cells. (**A-C**) IL-1α (**A**), Il-1β (**B**), and IL-18 (**C**) levels in the supernatants of cells infected with *S. flexneri* for the indicated times or following stimulation with 10 mM nigericin for 2 hours (without LPS priming). *n* = 3 experiments. Two-way ANOVA with Šidák *post hoc* test. Mean ± S.E.M. *, p<0.05; **, p<0.01; ***; p<0.001; ****, p<0.0001.

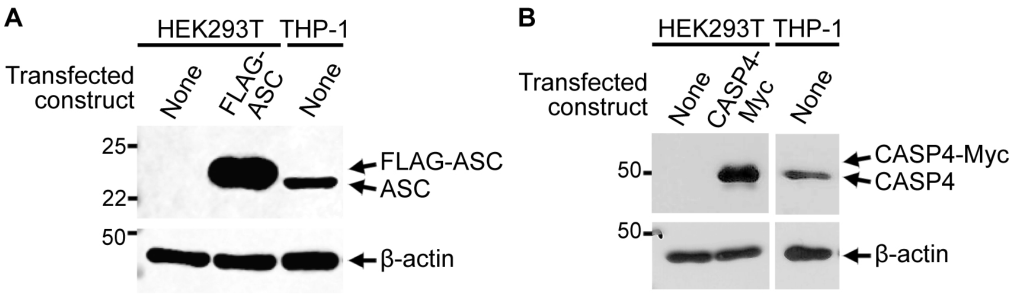


**Fig. S9. Absence of key innate immune proteins in HEK293T cells.**

Absence of detectable ASC and caspase-4 in HEK293T cells. Representative western blots; β-actin, loading control. (**A**) Levels of ASC in lysates of untransfected HEK293T cells. As controls, lysates of HEK293T cells transfected with FLAG-ASC and lysates of untransfected THP-1 cells are also shown. $n$ = 3 experiments. (**B**) Levels of caspase-4 (CASP4) in lysates of untransfected HEK293T cells. As controls, lysates of HEK293T cells transfected with CASP4-Myc and lysates of untransfected THP-1 cells are also shown. The two panels showing CASP4 are from the same blot, and the two panels showing β-actin are from the same blot. $n$ = >5 experiments. MW markers in kD.

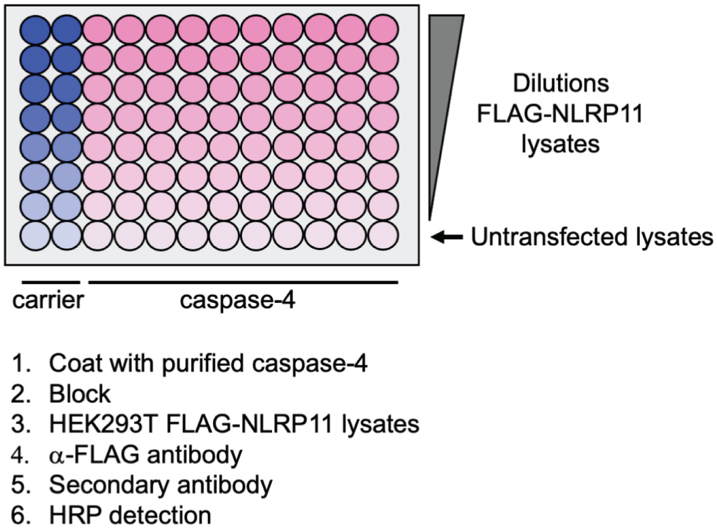


**Fig. S10. Plate binding assay for assessing interaction relationship of NLRP11 and caspase-4.**

Schematic of plate binding assay. Plate is coated with various concentrations of purified caspase-4 or carrier. After blocking, lysates of FLAG-NLRP11 transfected HEK293T cells are added at a range of dilutions, followed by antibody-based detection. Dilution of lysates of FLAG-NLRP11 transfected HEK293T cells was into lysates of untransfected HEK293T cells.

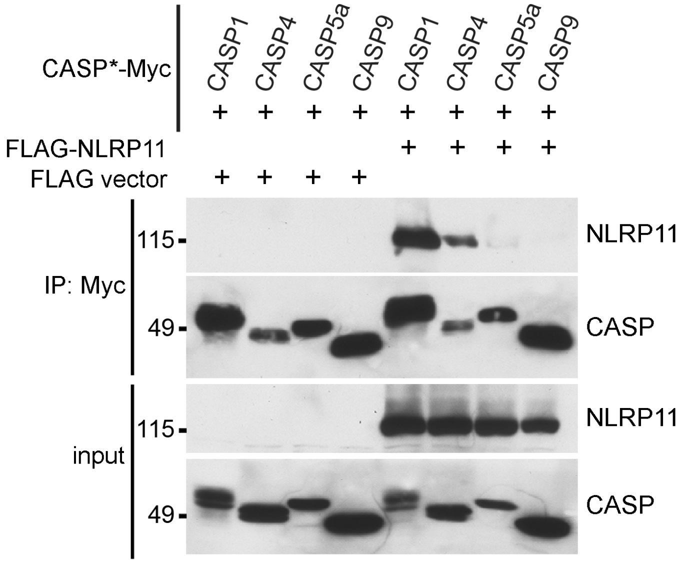


**Fig. S11. Specificity of NLRP11 interaction with caspases is independent of the tag.**

Specificity of NLRP11 for caspase-4 and caspase-1. Myc-tagged catalytically inactive caspases (CASP1[C285A]-Myc, CASP5a[C315A]-Myc, CASP9[C287S]-Myc) precipitation of FLAG-NLRP11. CASP*-Myc, catalytically inactive caspase. IP: Myc, precipitation with beads coated with Myc antibody. Representative western blots, detected with anti-FLAG (NLRP11) or anti-Myc antibody (caspases, CASP). *n* = 3 experiments. MW markers in kD.

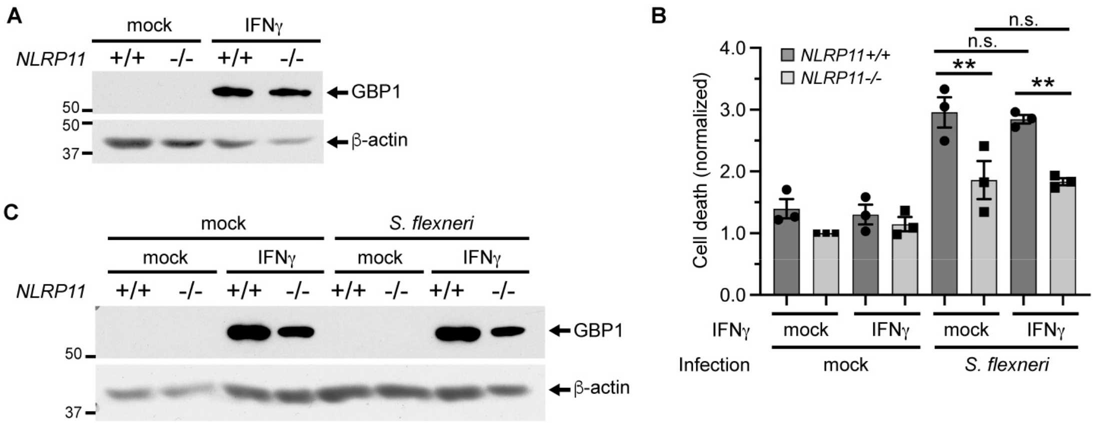


**Fig. S12. NLRP11 role in non-canonical inflammasome activation is independent of IFNγ and induction of GBP1.**

(**A**) Pre-treatment with IFNγ induced GBP1 similarly in NLRP11+/+ and NLRP11-/- THP-1 macrophages. Representative western blots of cell lysates. (**B**) Cell death during infection with *S. flexneri* depends on NLRP11, even following IFNγ induction of GBP1. (**C**) GBP1 is induced by pre-treatment with IFNγ but not by 2.5 hr of *S. flexneri* infection. Representative western blots of cell lysates from experiments presented in panel (**B**). $n$ = 3 experiments. MW markers in kD. Mean ± S.E.M. Two-way ANOVA with Tukey *post hoc* test. **, $p<0.01$; n.s., not significant.
